# Molecular basis of broad neutralization against SARS-CoV-2 variants including Omicron by a human antibody

**DOI:** 10.1101/2022.01.19.476892

**Authors:** Bin Ju, Qingbing Zheng, Huimin Guo, Qing Fan, Tingting Li, Shuo Song, Hui Sun, Senlin Shen, Xinrong Zhou, Lin Cheng, Wenhui Xue, Lingyan Cui, Bing Zhou, Xiangyang Ge, Haiyan Wang, Miao Wang, Shaowei Li, Ningshao Xia, Zheng Zhang

## Abstract

Omicron, a newly emerging SARS-CoV-2 variant, carried a large number of mutations in the spike protein leading to an unprecedented evasion from many neutralizing antibodies (nAbs). Here, we performed a head-to-head comparison of Omicron with other existing highly evasive variants in terms of their reduced sensitivities to antibodies, and found that Omicron variant is significantly more evasive than Beta and Mu variants. Of note, some key mutations occur in the conserved epitopes identified previously, especially in the binding sites of Class 4 nAbs, contributing to the increased Ab evasion. We also reported a broadly nAb (bnAb), VacW-209, which effectively neutralized all tested SARS-CoV-2 variants and even SARS-CoV. Finally, we determined six cryo-electron microscopy structures of VacW-209 complexed with the spike ectodomains of wild-type, Delta, Mu, C.1.2, Omicron, and SARS-CoV, and revealed the molecular basis of the broadly neutralizing activities of VacW-209 against SARS-CoV-2 variants. Overall, Omicron has once again raised the alarm over virus variation with significantly compromised neutralization. BnAbs targeting more conserved epitopes among variants will continue to play a key role in pandemic control and prevention.

**One sentence summary:** Structural and functional analyses reveal that a human antibody named VacW-209 confers broad neutralization against SARS-CoV-2 variants including Omicron by recognizing a highly conserved epitope.

## Main text

The coronavirus disease 2019 (COVID-19) caused by the infection of severe acute respiratory syndrome coronavirus 2 (SARS-CoV-2) is still a pandemic raging across the world. New variants, such as Alpha, Beta, Gamma, Delta, and Kappa, etc(*1–4*), keep emerging, causing a series of COVID-19 waves in different regions and countries worldwide. In addition to higher transmissibility and infectivity compared with the original wild-type (WT) virus, they could also escape, to a great extent, the neutralization of plasma and monoclonal neutralizing antibodies (nAbs) elicited by natural virus infection and vaccines(*2, 3, 5, 6*). For example, the Delta variant was first identified in India and has since become a long-time dominant variant rapidly spreading around the world due to its high transmissibility and some level of immune evasion by nAbs(*4, 7*). Its dominance can be revealed by the sequences collected on daily basis to the GISAID database since July 2021, more than 90% of which were Delta lineage (**Fig. 1A**). The Beta has long been regarded as the variant with the greatest reduction in neutralizing capacity(*8*). However, a recent study showed that Mu variant, once driving the epidemic in Colombia, was more evasive than Beta in the neutralizing activity(*9*). The Mu variant shared similar mutations in receptor binding domain (RBD) of spike with Beta, R346K/E484K/N501Y and K417N/E484K/N501Y, respectively. After that, the C.1.2 variant was first identified and mainly reported in South Africa, also carrying three mutations (Y449H/E484K/N501Y) in RBD, which was similar to Beta(*10*). Although Beta, Mu, and C.1.2 variants did not give rise to a global pandemic like Delta (**Fig. 1, A and B**), their occurrence were warning of possible variants with even more serious escape from the neutralization of pre-existing antibodies.

**Figure 1.**
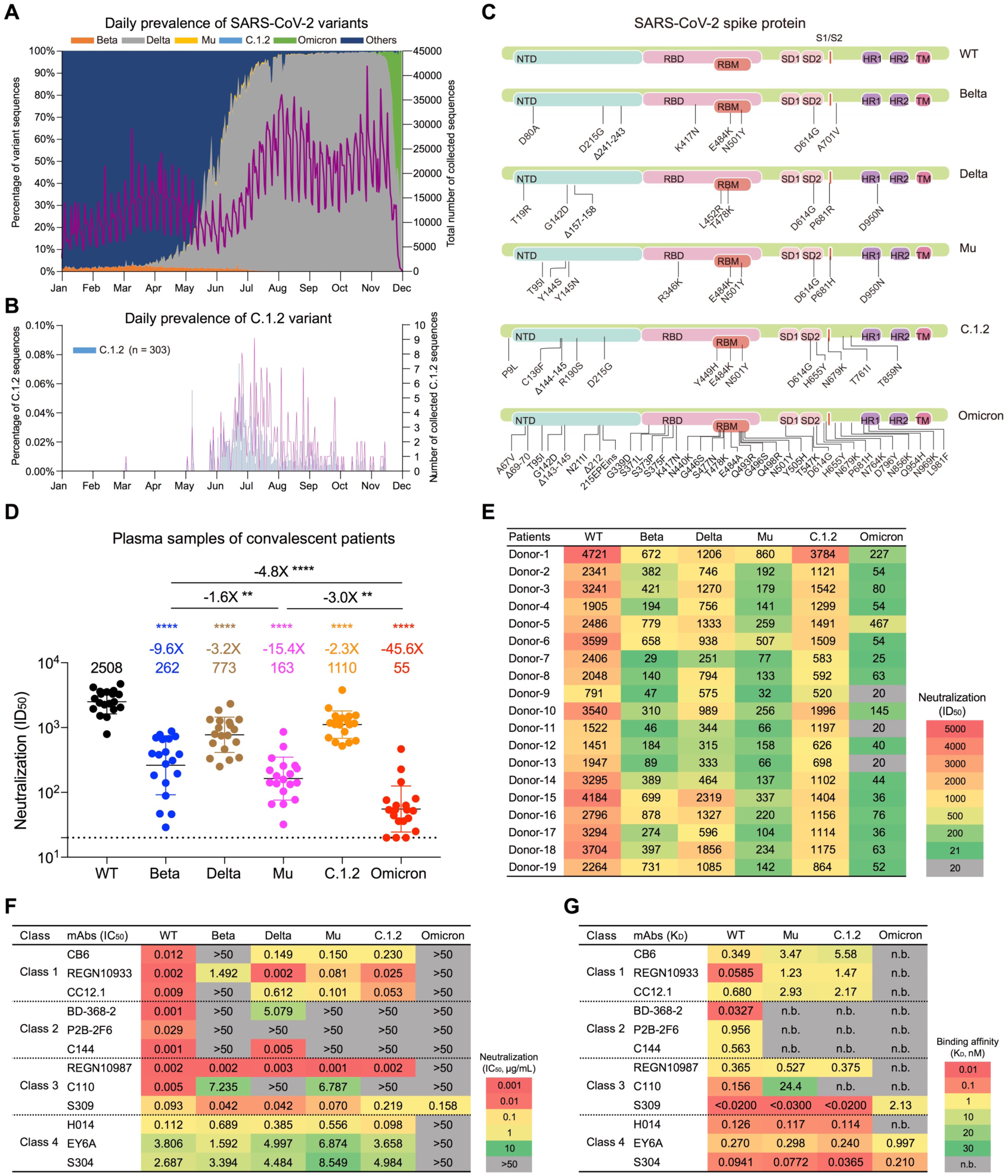
Resistance of SARS-CoV-2 variants including Omicron to the neutralization by convalescent plasma and monoclonal nAbs. **(A)** Daily prevalence of the variants and their distributions in all collected SARS-CoV-2 sequences in the world deposited to GISAID in 2021. The left y-axis indicates the percentage of each variant collected everyday, which is displayed by areas in different colors. Orange: Beta, Gray: Delta, Yellow: Mu, Light blue: C.1.2, Green: Omicron, Dark blue: Others. The right y-axis indicates the total number of collected sequences each day, which is displayed by the purple line. **(B)** The prevalence of C.1.2 variant was separated to shown due to its rarity among the reported sequences. **(C)** The landscape of key mutations in spike proteins of SARS-CoV-2 variants used in this study. The full-length mutated spike genes were synthesized to construct pseudoviruses. The WT, Beta, Delta, Mu, C.1.2, and Omicron variants were shown from top to bottom. **(D)** The neutralization of WT SARS-CoV-2 and variants by plasma samples of 19 convalescent patients infected with the WT virus and recovered from the first wave of COVID-19 pandemic. The GMT of nAbs, fold change, and statistical analysis were calculated and noted on the top of each column. The data are shown in Geometric mean ± SD. “-” indicates decreased neutralization activity. Fold changes in GMTs were compared between each variant and WT or between two variants. Statistical analysis was performed with a paired *t* test using GraphPad Prism 9 software. **: P < 0.01, ****: P < 0.0001. The horizontal dashed line indicates the limit of detection (1:20 dilution) for the neutralizing assay. The non-neutralizing data below the limit were set to 20 for visualization. **(E)** The neutralization of each plasma sample against WT and variants was represented in ID_50_ value. **(F)** The neutralization (IC_50_) of 12 representative nAbs of Class 1 to 4 against WT SARS-CoV-2 and variants. The cutoff value of neutralization was set as 50 μg/mL. **(G)** The binding affinity (K_D_) of 12 representative nAbs to RBD proteins of WT SARS-CoV-2 and variants by SPR. The neutralizing potency or binding affinity is highlighted in different colors. Red: high, Yellow: moderate, Green: weak, Gray: non-neutralizing or not binding (n.b.). The data are means of two independent experiments.

A new variant, Omicron, was first detected in South Africa in November 2021, and got dominant in many regions where Delta is prevalent with the short doubling time of cases(*11*). Omicron has been classified as a variant of concern (VOC) by the World Health Organization (WHO), whose spike protein carried more than 30 mutations including amino acid substitutions, deletions, and insertions(*12*). So many mutations, especially in the RBD (15 substitutions), had never appeared in any previous variants, suggesting that Omicron may sharply escape the nAbs elicited by the WT virus or vaccine. Some recent studies indeed showed that Omicron significantly reduced the neutralization of plasma from convalescent patients or individuals immunized with several major SARS-CoV-2 vaccines. The Omicron variant also largely escaped the neutralization of monoclonal nAbs that have been approved for Emergency-Use-Administration (EUA), decreasing or abolishing their neutralizing activities(*11–14*). However, the molecular basis of the enhanced antibody evasion is not well defined, especially for comparison of the Omicron to Beta, Delta, Mu, and C.1.2 variants with more serious escape from the nAbs. In addition, it lacks a head-to-head comparison of the structural and functional characterization for the novel broadly nAb (bnAb) against these variants and even SARS-CoV.

We previously developed and validated a pseudovirus neutralization assay to evaluate the neutralizing activities of plasma and mAbs against the WT SARS-CoV-2 and variants(*15–17*). On top of it, we constructed Mu, C.1.2, and Omicron pseudoviruses (**Fig. 1C**). To make a comprehensive evaluation of the susceptibility of Omicron, we summarized plasma samples from 19 individuals who were infected with the WT virus and had recovered from COVID-19 in the first wave of pandemic, and then measured and compared their neutralizing activities against the WT, Beta, Delta, Mu, C.1.2, and Omicron variants, head to head (**Fig. 1D and fig. S1**). In consistence with a previous study(*9*), the Mu variant showed more serious resistance to nAbs than Beta (15.4-fold vs. 9.6- fold). In contrast, the Delta and C.1.2 variants showed the relatively weaker escape from the neutralization of pre-existing antibodies (3.2-fold and 2.3-fold, respectively). Remarkably, we found that the geometric mean titer (GMT) of nAbs against Omicron decreased, three folds greater than Mu (55 vs. 163), and over 40 folds compared with that against WT (GTM = 2508). More seriously, some plasma (3/19) lost their neutralizing activities against Omicron, whose 50% inhibitory dilution (ID_50_) was less than the dilution of 1:20 (**Fig. 1E**). Overall, the largely enhanced antibody evasion by Omicron variant is unprecedented, and the appearance of Omicron is another warn of virus escape.

To study the mechanism of antibody escape, we summarized and prepared a panel of 12 published nAbs binding to RBD of SARS-CoV-2 with clear structural information. The RBD recognizes the cellular receptor (angiotensin-converting enzyme 2, ACE2) and mediates the virus entry into the host cell, which is the primary target for blocking viral infection(*18, 19*). Therefore, RBD-specific nAbs contribute a lot to the neutralization of SARS-CoV-2. Here, we used these 12 nAbs recognizing diverse epitopes to mimic the polyclonal antibodies in plasma to explore what kind of nAbs were mostly affected by the mutations of variants and the molecular basis of antibody evasion. In general, the nAbs can be classified into classes 1, 2, 3, and 4 based on the competition with ACE2 and the binding model to RBD (**fig. S4A**)(*20, 21*). The Beta mainly escaped from the neutralization of nAbs of Class 1 and 2, yet Delta, Mu, and C.1.2 partially or totally escaped the nAbs of Class 2 and 3 we tested. Notely, the neutralizing and binding activities of Class 4 antibodies were not affected by these variants (**Fig. 1, F and G; fig. S2 and S3**). However, Omicron abolished the neutralization of nearly all nAbs we tested (11/12, except S309) across Class 1 to 4 (**Fig. 1F**), explaining why Omicron has the most serious antibody evasion so far.

The structural analysis further showed that some key mutations were located in or near the footprint of nAbs on the RBD (**fig. S4A**). The K417N and Q493R substitutions mainly affect the recognition of Class 1 nAbs to Omicron RBD. For example, the overall binding mode of CB6 had been changed. CB6 interacts to the K417 residue on RBD with Y33, Y52, and D104 residues of heavy chain through hydrogen bond and salt bridge interactions. Q493 is another key epitope residue for Class 1 nAbs binding to RBD, which make a salt bridge with heavy-chain Y109 residue (**fig. S4B**). E484 on WT RBD forms hydrogen bond and salt bridge interactions with R100, R102, and D106 of BD-368-2 heavy chain. Therefore, mutations at E484 usually result in complete insensitivity of Class 2 nAbs (**fig. S4C**). The mechanisms of Class 1 and 2 nAbs escape mentioned above have been stated on Beta, Gamma, and Kappa variants in previous reports(*2, 5, 22-24*). For REGN10987, an antibody of Class 3, the G446S mutation may diminish the binding of this class of nAbs to RBD (**fig. S4D**). As mentioned above, the Omicron was the first variant to escape Class 4 nAbs. The structural analysis of H014 showed that S371L may mediate the resistance of Omicron to this class of nAbs (**fig. S4E**). Previously, REGN10987-like nAbs and Class 4 nAbs were considered to have the most broadly neutralizing activities, whose targeting epitopes were relatively conserved among various SARS-CoV-2 variants(*4, 5, 7, 25, 26*). G446S and S371L are both newly identified signature mutations of the Omicron, which confer the great antibody resistance and have changed our understanding about the bnAbs. Fortunately, some minority of existing nAbs are effective to Omicron. Consistent with previous studies(*11, 27*), S309 retains a strong binding affinity to Omicron-RBD and an effective neutralizing activity (**Fig. 1, F and G**), despite the existence of G339 and N440 residues as part of its epitopes (**fig. S4D**).

Then we explore other bnAbs against Omicron, especially those bind to distinct epitopes away from S309. Previously, we identified 9 monoclonal nAbs from 2 individuals immunized with the SARS-CoV-2 inactivated vaccine and measured their cross-neutralizing activities against Kappa and Delta(*17*). Here, we further evaluated the neutralization of these nAbs against other important variants including Alpha, Beta, Gamma, Mu, especially Omicron, and so on (**Fig. 2A and fig. S5**). Their neutralizing breadths ranged from 54% to 100% in the tested 13 SARS-CoV-2 pseudoviruses, and a few of the nAbs (3/9) could still neutralize Omicron. VacW-92 and VacW-120 binding to overlapped epitopes with S309(*17*) effectively neutralized Omicron (1.246 μg/mL and 0.273 μg/mL, respectively) but not Delta. VacW-209 could neutralize all variants tested here with a high potency (Geometric IC_50_ = 0.063 μg/mL). Meanwhile, the binding affinities of VacW-209 to Mu, C.1.2, and Omicron RBD proteins may contribute to its broadly neutralizing activity (**Fig. 2B and fig. S6**). SARS-CoV is closely related to SARS-CoV-2 sharing 80% of amino acid sequence identity in their spike proteins(*28*). Therefore, we also detected the cross-reaction of VacW-209 to SARS-CoV, which displayed both highly neutralizing activity (0.141 μg/mL) and binding affinity (0.540 nM) (**Fig. 2, C and D**). VacW-209 strongly competed with ACE2 for binding to RBD, revealing its neutralizing mechanism of high potency (**Fig. 2E**). Then, we measured the competition of VacW-209 with 15 typical nAbs of Class 1 to 4, which revealed that VacW-209 bound to an epitope overlapped with that of both Class 1 and Class 4 nAbs (**Fig. 2F**). It was found that VacW-209 did not compete with two approved nAb drugs (REGN10987 and S309), suggesting that it could be used both independently and in combination with these nAbs. We therefore evaluated the neutralizing activities of VacW-209+REGN10987 and VacW-209+S309 against WT, Beta, Delta, and Omicron,respectively (**Fig. 2, G and H**). REGN10987 completely lost its neutralization against Omicron, but it could be effectively rescued through the combination with VacW-209. Considering that S309 and VacW-209 are both potent nAbs against all identified SARS-CoV-2 variants so far and SARS-CoV, the combination will open up the way against virus escape in the future.

**Figure 2.**
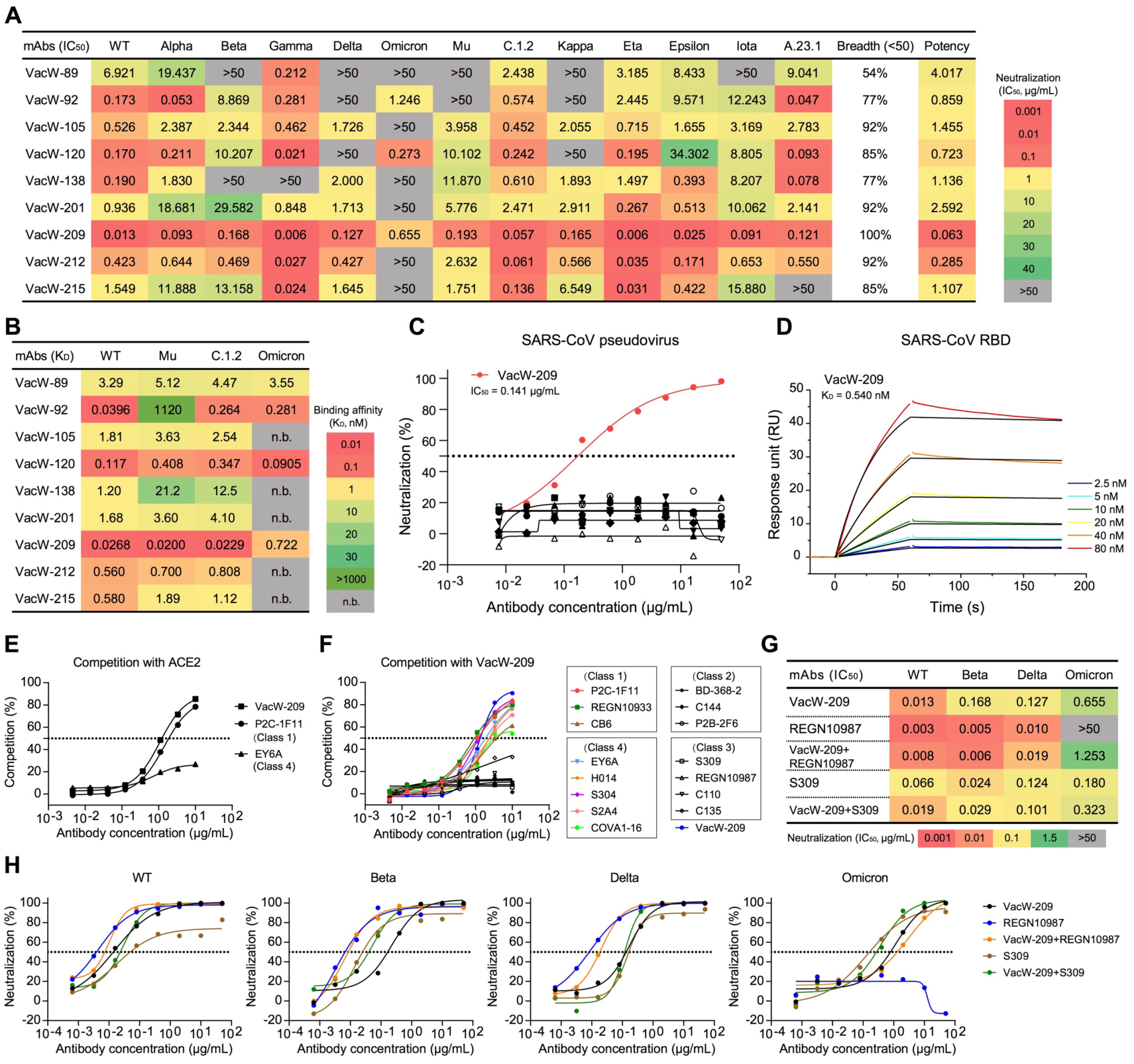
Identification of bnAbs against SARS-CoV-2 variants including Omicron. **(A)** The neutralization (IC_50_) of 9 nAbs isolated from individuals vaccinated with WT SARS-CoV-2 vaccine against 13 tested pseudoviruses. The cutoff value of neutralization was set as 50 μg/mL. The neutralizing breadth was calculated as the percentage of viruses neutralized by each nAb. Geometric mean potency was calculated by the neutralizing value of less than 50 μg/mL. **(B)** The binding affinity (K_D_) of 9 nAbs to RBD proteins of WT SARS-CoV-2 and variants by SPR. The neutralizing and/or binding activities of 9 nAbs to WT, Delta, and Kappa have been reported in our previous study(*17*), which are re-tested here for a head-to-head comparison with other variants. **(C)** The neutralizing activities of 9 nAbs against SARS-CoV. VacW-209 is marked in red, the other non-neutralizing mAbs are marked in black. **(D)** The binding affinity of VacW-209 to SARS-CoV RBD by SPR. **(E)** Competition ELISA of VacW-209 with human ACE2 for binding to WT RBD. P2C-1F11 was a known competitive antibody as a positive control. EY6A was a known non-competitive antibody as a negative control. **(F)** Competition ELISA of VacW-209 with 15 representatives nAbs of four classes and with itself for binding to WT RBD. The neutralizing activities of VacW-209 combined with REGN10987 or S309 against WT, Beta, Delta, and Omicron were represented in IC_50_ values **(G)** and curves **(H)**. The data represented here are means of at least two independent experiments. The neutralizing potency or binding affinity is highlighted in different colors. Red: high, Yellow: moderate, Green: weak, Gray: non-neutralizing or not binding (n.b.). The curve of neutralization, binding affinity, or competition was one out of similar results. A cutoff value of 50% is indicated by a horizontal dashed line in neutralization and competition.

To define the structural basis of the broadly neutralizing activity of VacW-209, we resolved the cryo-electron microscopy (cryo-EM) structures of the antigen-binding fragment (Fab) of VacW-209 in complex with the spike proteins of SARS-CoV-2 WT, Delta, Mu, C.1.2, Omicron, and SARS-CoV, respectively. Six cryo-EM structures of trimeric spike based immune complexes at 2.98-3.45 Å revealed nearly identical binding modes of VacW-209 (**Fig. 3, A to F; fig. S7 to S12; and table S1**). Three VacW-209 Fabs bound to a completely opened S with three “up” RBDs. We then performed the focus refinement of regions of Fab-bound RBDs of WT, Delta, Mu, C.1.2, and Omicron, while the Fab-bound SARS-CoV RBD was weakly resolved due to structural flexibility (**Fig. 3, G to K**). High-resolution structures revealed that the binding epitope of VacW-209 completely evaded the key RBD mutations in variants of Delta, Mu, C.1.2, and rarely overlapped with mutations in Omicron (**Fig. 3, H to K, lower**). The footprints of VacW-209 on WT-RBD and Omicron-RBD were slightly different, and three mutations in Omicron (K417N, S373P, and S375F) were involved in the nAb-RBD interaction (**Fig. 3, L and M**).

**Figure 3.**
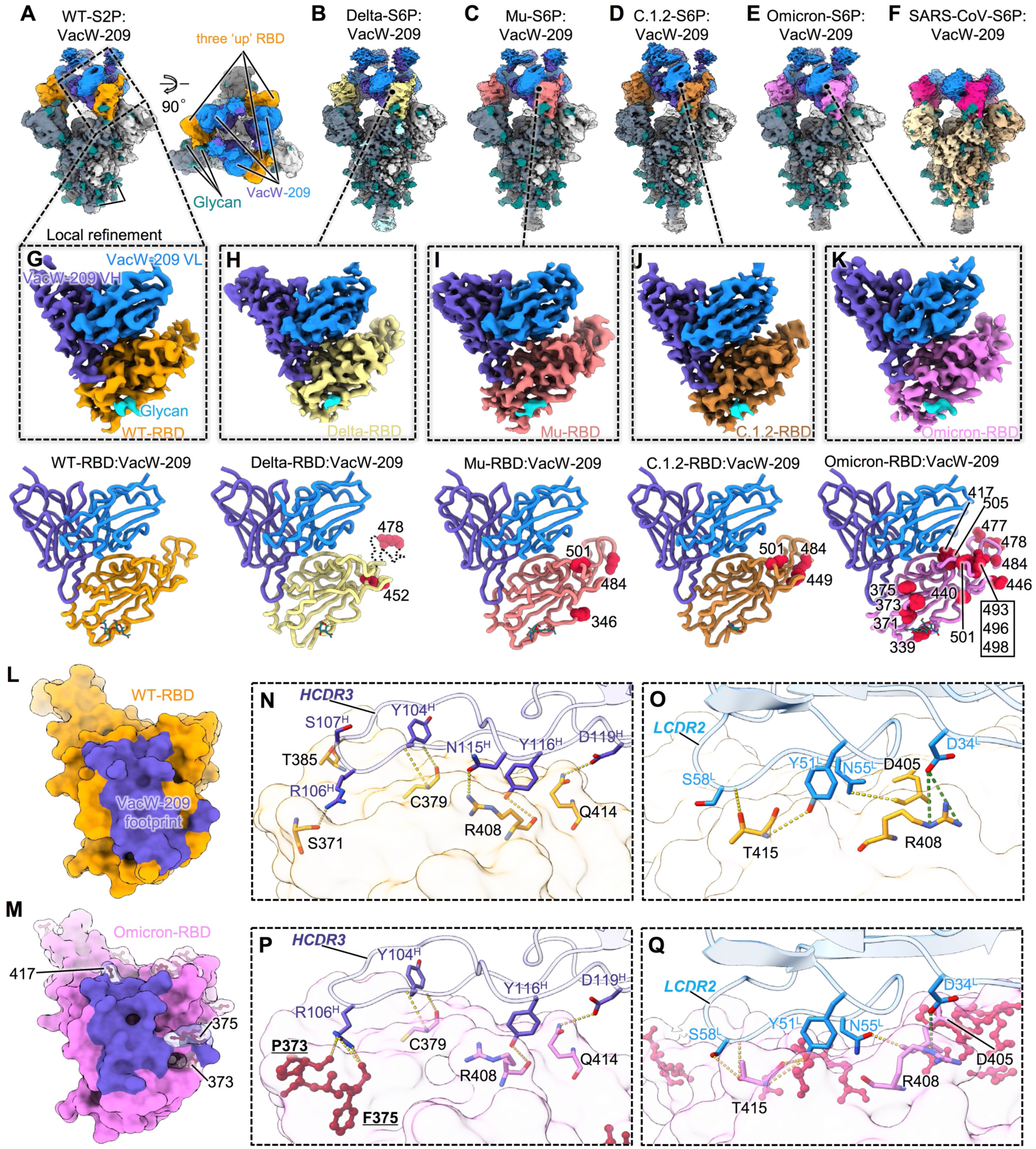
Cryo-EM structures of VacW-209 complexed with spike proteins of WT SARS-CoV-2, variants, and SARS-CoV. **(A-F)** Cryo-EM density maps of VacW-209 in complex with spike proteins of WT-S2P (A), Delta-S6P (B), Mu-S6P (C), C.1.2-S6P (D), Omicron-S6P (E), and SARS-CoV-S6P (F). **(G-K)** Cryo-EM density maps (upper) and corresponding atomic models (lower) of local refinement of VacW-209 in complex with WT-RBD (G), Delta-RBD (H), Mu-RBD (I), C.1.2-RBD (J), and Omicron-RBD (K). Models are represented as cartoon and key mutations on RBD are highlighted as red balls. **(L, M)** The binding footprints of VacW-209 (colored in purple) on WT-RBD (orange surface representation) (L) and Omicron-RBD (pink surface representation) (M). The mutated residues are rendered as red sticks with transparent surface representation on Omicron RBD and those involved in VacW-209 interactions are labeled. **(N, O)** Interaction details between WT-RBD and VacW-209 heavy chain (N) and light chain (O). **(P, Q)** Interaction details between Omicron-RBD and VacW-209 heavy chain (P) and light chain (Q). Hydrogen bonds and salt-bridges are labeled as orange and dark green dotted lines respectively.

We next analyzed the interaction details of VacW-209 binding to WT and Omicron spikes, respectively, and revealed that VacW-209 mainly used its extremely long heavy loop at complementarity determining region (CDR) 3 to mediate spike protein recognition. Besides, the light chain CDR2 and D34 from LCDR1 were also involved in nAb-RBD interactions (**Fig. 3, N to Q**). For WT-RBD, residues 371, 379, 408, 414, and 415 formed a total of 11 hydrogen bonds and 2 salt bridges to VacW-209 and created an interaction network between them (**Fig. 3, N and O**). The heavy chain R106 inserted its long side chain into the pocket formed by RBD aa. 371-385, which contained three key mutations of Omicron (S371L/S373P/S375F) (**fig. S13, A to C**). Although VacW-209 showed a decreased neutralization against Omicron compared with that against WT, to some extent (**Fig. 2A**), our structural analysis showed that the mutations surrounding RBD aa. 371-385 loop seemed not to obviously affect the binding of VacW-209 since the S373P and S375F built three new hydrogen bonds with heavy chain R106 (**Fig. 3P**). Other Omicron mutations were not involved in the hydrogen bond interactions, and there were a total of 12 hydrogen bonds and 1 salt bridges formed (**Fig. 3, P and Q**), which were comparable in total to that in WT. We further found that the binding of VacW-209 to Omicron RBD need a slight conformational change of 371-385 loop containing S371L/S373P/S375F mutations (**fig. S13, D and E**), which may partly account for the reduced neutralization of VacW-209 against Omicron variant.

Finally, we compared the binding mode of VacW-209 to several nAbs of Class 1 to 4 and defined a new binding mode of VacW-209 which bound to an epitope between Class 1 and Class 4, yet not overlapping with that of Class 2 or Class 3 (**Fig. 4A**). Despite some minor differences in details, the binding of VacW-209 to RBDs of SARS-CoV-2 WT, Delta, Mu, C.1.2, and Omicron were all mediated by the long HCDR3 (in particular R106, Y116, and D119), LCDR2 (in particular Y51, N55, and S58), and LCDR1 residue D34 (**Fig. 4, B to F**). We also explored the potential binding sites of VacW-209 on other variant RBDs as well as SARS-CoV RBD based on the binding characterization revealed in the WT-S2P:VacW-209 (**Fig. 4, G to K**). The epitope of VacW-209 nearly excluded all of above RBD mutations and was highly conserved between SARS-CoV-2 and SARS-CoV (**Fig. 4, C to K**). The epitope was mainly comprised of RBD aa. 376-385 and 405-416, which was highly conserved among SARS-CoV-2 variants and there were only three amino acid substitutions between SARS-CoV-2 and SARS-CoV (**Fig. 4L**).

**Figure 4.**
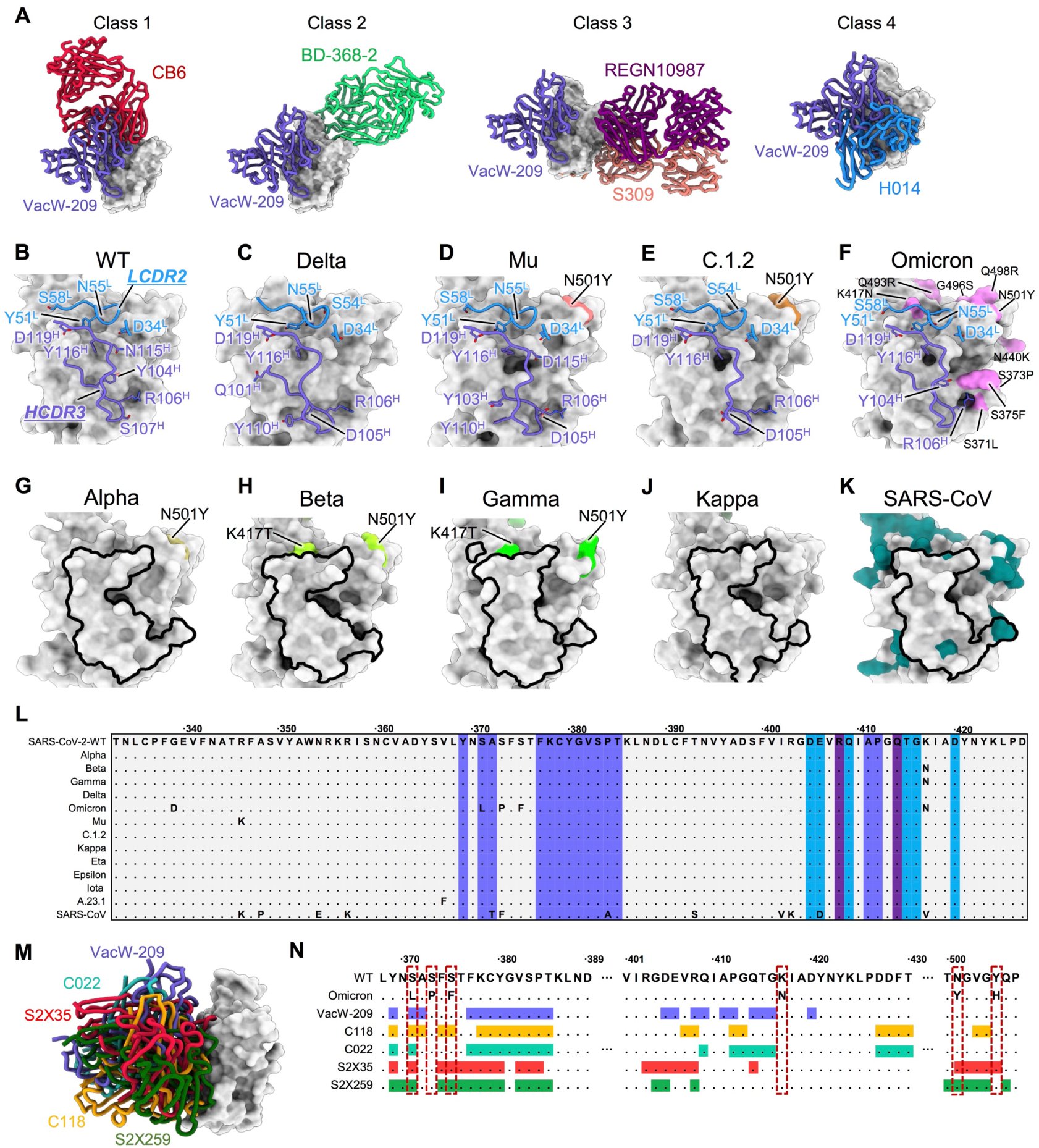
Binding mode and the epitope conservation of VacW-209. **(A)** Comparison of the binding mode of VacW-209 to representative Class 1-4 nAbs. Class 1: CB6 (7C01), Class 2: BD-368-2 (7CHH), Class 3: REGN10987 (6XDG) and S309 (7R6W), and Class 4: H014 (7CAH). **(B-F)** Structural compassion of VacW-209 bound to SARS-CoV-2 WT-RBD (B), Delta-RBD (C), Mu-RBD (D), C.1.2-RBD (E), and Omicron-RBD (F). **(G-K)** Structure comparison of RBDs of Alpha (G) (7LWV), Beta (H) (7VX1), Gamma (I) (7M8K), Kappa (J) (7VXE) variants, and SARS-CoV (K) (7JN5). RBDs are shown as gray surface representations and key mutations on RBDs are labeled. The modeled VacW-209 footprints are shown on (G-K) based on the epitope information revealed in SARS-CoV-2 WT-S. **(L)** RBD sequences of SARS-CoV-2 WT and its 12 variants as well as SARS-CoV with highlighted footprint of VacW-209 (slate blue: heavy chain, dodger blue: light chain, dark blue: both chains). **(M)** VacW-209-like nAbs and their binding modes on RBD. VacW-209, C118 (7RKS), C022 (7RKU), S2X35 (7R6W), and S2X259 (7M7W) are shown as sticks and colored in blue, orange, cyan, red, and green, respectively. **(N)** RBD sequence of SARS-CoV-2 WT and Omicron variant with highlighted footprints of VacW-209, C118, C022, S2X35, and S2X259. Amino acids substitutions revealed on Omicron variant are boxed.

The similar binding mode of VacW-209 was also found in some previously reported nAbs including C118, C022, S2X35, and S2X259(*29–31*) (**Fig. 4M**). Available structural information revealed that the aforementioned four nAbs and VacW-209 shared lots of epitope residues located in conserved RBD aa. 376-385 and 405-416, while with diverse coverage of key mutations of Omicron (**Fig. 4N; fig. S14, A to E**). Of these mutations, S371L, S373P, S375F, K417N, N501Y, and Y505H were structurally close to or involved in the binding epitopes of VacW-209-like nAbs. In the head-to-head comparison, although C022 and S2X35 showed significant reductions of neutralization against Omicron, these VacW-209-like nAbs generally maintained effectively neutralizing and binding activities to various SARS-CoV-2 variants and even SARS-CoV (**fig. S14, F and G; fig. S15**). The molecular mechanism underlying why these similar nAbs display diverse neutralizing activities need to be elucidated in the future.

In conclusion, the bnAb described here, VacW-209, idenfies a highly conserved epitope on the RBDs among SARS-CoV-2 variants overlapping with the ACE2- binding site, responsible for its potent neutralization. VacW-209 could strongly compete with Class 4 nAbs, indicating the potential cross-neutralization against sarbecoviruses. These VacW-209-like nAbs shared a similar antibody response to both SARS-CoV-2 and SARS-CoV, highlighting a candidate target for the universal vaccine design. As a promising antibody therapeutics, VacW-209, alone or in combination with S309, could also be used as countmeasure against SARS-CoV-2 variants including Omicron and even other forthcoming sarbecoviruses in the future.

## Acknowledgments

We thank all participants who recovered from the COVID-19 and who received SARS-CoV-2 inactivated vaccines. We also thank all of the healthy providers from Shenzhen Third People’s Hospital for the work they have done. This study was supported by the National Science Fund for Distinguished Young Scholars (82025022), the National Natural Science Foundation of China (92169204, 82002140, 82171752, 82101861, 81991491), the National Key Plan for Scientific Research and Development of China (2021YFC0864500), the Guangdong Basic and Applied Basic Research Foundation (2021B1515020034, 2019A1515011197, 2021A1515011009, 2020A1515110656), the Shenzhen Science and Technology Program (RCYX20200714114700046), and the Science and Technology Innovation Committee of Shenzhen Municipality (JSGG20200207155251653, JSGG20200807171401008, KQTD20200909113758004, JCYJ20190809115617365, SGG20210901145200002).

## Author contributions

Z.Z. is the principal investigator of this study. Z.Z., N.X., S.L., and B.J. conceived and designed the study. B.J., Q.Z., H.G., Q.F., T.L., S. Song, and H.S. performed all experiments and analyzed data together with S. Shen, X.Z., L.Cheng, W.X., L.Cui, B.Z., X.G., H.W., and M.W.. Z.Z., N.X., S.L., B.J., and Q.Z. participated in discussion of the results and wrote the manuscript. All authors read and approved this version of manuscript.

## Data and materials availability

We are happy to share reagents and information in this study upon request. Structure coordinates are deposited in the Protein Data Bank under accession codes XXXX (WT-S2P:VacW-209), XXXX (Delta-S6P:VacW-209) XXXX (Mu-S6P:VacW-209), XXXX (C.1.2-S6P:VacW-209), and XXXX (Omicron-S6P:VacW-209). The corresponding EM density maps have been deposited in the Protein Data Bank under accession numbers XXXX (WT-S2P:VacW-209), XXXX (Delta-S6P:VacW-209) XXXX (Mu-S6P:VacW-209), XXXX (C.1.2-S6P:VacW-209), XXXX (Omicron-S6P:VacW-209), and XXXX (SARS-CoV-S6P:VacW-209).

## Supplementary information

### Materials and Methods

#### Study approval and plasma samples

This study was approved by the Ethics Committee of Shenzhen Third People’s Hospital, China (approval number: 2020-084). All participants had provided written informed consent for sample collection and subsequent analysis. All plasma samples were collected from 19 convalescent patients at Month 6 post recovery from the early COVID-19 pandemic. All plasma samples were stored at -80 °C in the Biobank of Shenzhen Third People’s Hospital and heat-inactivated at 56 °C for 1 h before use.

#### Expression and purification of monoclonal neutralizing antibodies

Gene sequences of published neutralizing antibodies (nAbs) downloaded from the National Center of Biotechnology Information (NCBI) were synthesized and cloned into the human full-length IgG1 expression vectors by Sangon Biotech and GenScript. Their protein data bank (PDB) codes were CB6 (7C01), REGN10933 (6XDG), CC12.1 (6XC2), BD-368-2 (7CHH), P2B-2F6 (7BWJ), C144 (7K90), REGN10987 (6XDG), C110 (7K8V), S309 (6WPS), H014 (7CAI), EY6A (6ZCZ), S304 (7JW0), P2C-1F11 (7CDI), C135 (7K8Z), S2A4 (7JVC), COVA1-16 (7JMV), C118 (7RKS), C022 (7RKU), S2X35 (7R6W), and S2X259 (7RAL), respectively. Paired heavy-chain and light-chain plasmids were co-transfected into 293 F cells. Monoclonal antibodies (mAbs) were purified from cell supernatants after five days using protein A columns according to the manufacturer’s instructions (National Engineering Research Center for Biotechnology). Purified antibodies were quantified by NanoDrop and stored at 4 °C before use.

Nine monoclonal nAbs (VacW-89, VacW-92, VacW-105, VacW-120, VacW-138, VacW-201, VacW-209, VacW-212, and VacW-215) were isolated from 2 individuals who received two doses of SARS-CoV-2 inactivated vaccines (WIBP-CorV, the Sinopharm COVID-19 vaccine, Wuhan Institute of Biological Products Co., Ltd). Their cross-neutralization of SARS-CoV-2 Kappa and Delta variants have been reported in our previous study(*1*). Here, we further tested their neutralizing activities against Beta, Mu, Omicron, and other variants.

#### Incidence analysis of SARS-CoV-2 variants

Daily numbers of collected SARS-CoV-2 sequences in the world in 2021 were obtained from the GISAID (https://www.gisaid.org/) including Beta, Delta, Mu, C.1.2, Omicron, and other variants. Graphs were plotted using GraphPad Prism 9 software.

#### SARS-CoV-2 and SARS-CoV pseudoviruses

Pseudovirus was generated by co-transfection of HEK-293T cells with a spike-expressing plasmid and an env-deficient HIV-1 backbone vector (pNL4-3.Luc.R-E-) as previously described(*1–3*). Two days post co-transfection, the culture supernatant was harvested, clarified by centrifugation, filtered, and stored at -80 °C. The infectious titer was measured by luciferase activity in the HEK-293T-hACE2 cells using Bright-Lite Luciferase reagent (Vazyme Biotech). Detailed sequence information of spike protein was listed below, respectively.

<u>SARS-CoV-2 wild-type (WT)</u>:

Wuhan-Hu-1, accession number: NC_045512;

<u>SARS-CoV-2 Beta:</u>

D80A, D251G, 242-243del, K417N, E484K, N501Y, D614G, A701V;

<u>SARS-CoV-2 Delta:</u>

T19R, G142D, 157-158del, L452R, T478K, D614G, P681R, D950N;

<u>SARS-CoV-2 Mu:</u>

T95I, Y144S, Y145N, R346K, E484K, N501Y, D614G, P681H, D950N;

<u>SARS-CoV-2 C.1.2:</u>

P9L, C136F, 144-145del, R190S, D215G, Y449H, E484K, N501Y, D614G, H655Y, N679K, T761I, N859N;

<u>SARS-CoV-2 Omicron:</u>

A67V, 69-70del, T95I, G142D, 143-145del, N211I, 212del, 215EPEins, G339D, S371L, S373P, S375F, K417N, N440K, G446S, S477N, T478K, E484A, Q493R, G496S, Q498R, N501Y, Y505H, T547K, D614G, H655Y, N679K, P681H, N764K, D796Y, N856K, Q954H, N969K, L981F;

<u>SARS-CoV wild-type (WT):</u>

CUHK-W1, accession number: AAP13567.1.

### Recombinant receptor binding domain (RBD) and trimeric spike proteins

The SARS-CoV-2 WT RBD protein (residues 319-541) was expressed with a His tag at the C-terminus. The RBD expression vectors of Mu and C.1.2 were constructed by point mutation based on the pCMV3-WT-RBD-His (Sino Biological, VG40592-CH). These RBD plasmids were transiently transfected into 293 F cells, respectively. After 5 days, RBD proteins were collected from the supernatant using Ni-sepharose fast-flow 6 resin (GE Healthcare) and eluted by 250 mM imidazole. The RBD proteins of SARS-CoV-2 Omicron (40592-V08H121) and SARS-CoV (40150-V08B2) were purchased from Sino Biological. All RBD proteins were used to measure the binding affinity of mAbs. The SARS-CoV-2 WT S-2P protein (residues 1-1208) was prepared in our previous study(*4, 5*), carrying two stabilizing Pro substitutions (986 and 987) and ‘‘GSAS’’ substitutions at the furin cleavage site (682-685). The other S-6P proteins of SARS-CoV-2 Delta, Mu, C.1.2, and SARS-CoV were expressed and purified in this study. Compared with S-2P, S-6P version contained additional four Pro substitutions (F817P, A892P, A899P, and A942P). The coding genes of spike ectodomain followed by a foldon trimerization motif, a His tag, and a flag tag at the C-terminus were synthesized and cloned into the pcDNA3.4 vector by GenScript. All S-6P plasmids were transiently transfected into 293 F cells, respectively. After 5 days, the secreted S-6P proteins were purified from the supernatant with Ni-sepharose fast-flow 6 resin (GE Healthcare) and 250 mM imidazole. The SARS-CoV-2 Omicron S-6P protein (40589-V08H26) was purchased from Sino Biological. All S-2P or S-6P proteins mentioned here were further used to perform the cryo-electron microscopy (Cryo-EM) experiment.

### Pseudovirus-based neutralizing assay

To determine the neutralizing activity, plasma samples or purified mAbs were serially diluted and then incubated with an equal volume of pseudovirus at 37 °C for 1 h. HEK-293T-hACE2 cells were subsequently added to the 96-well plates. After a 48-h incubation, the culture medium was removed, and 100 μL of Bright-Lite Luciferase reagent was added to the cells. After an incubation at RT for 2 mins, 90 μL of cell lysate was transferred to the 96-well white solid plates for measurements of luminescence using Varioskan™ LUX multimode microplate reader (Thermo Fisher Scientific). The 50% inhibitory dilution (ID_50_) for plasma or 50% inhibitory concentration (IC_50_) for mAbs was calculated using GraphPad Prism 9 software by log (inhibitor) vs. normalized response - Variable slope (four parameters) model.

### Binding analysis by surface plasmon resonance (SPR)

The binding assays of mAbs to the RBD proteins were performed using the Biacore 8K system (GE Healthcare). Specifically, one flow cell of the CM5 sensor chips were covalently coated with the RBD proteins in 10 mM sodium acetate buffer (pH 5.0) for a final RU (response units) around 250, whereas the other flow cell was left uncoated and blocked as a control. All the assays were run at a flow rate of 30 µL/min in HBS-EP buffer (10 mM HEPES pH 7.4, 150 mM NaCl, 3 mM EDTA, and 0.05% Tween-20). Serially diluted mAbs were injected for 60 s respectively and the resulting data were fit in a 1:1 binding model with Biacore Evaluation software (GE Healthcare). Every measurement was performed two times and the individual values were used to produce the mean affinity constant.

### Competition enzyme linked immunosorbent assay (ELISA)

The SARS-CoV-2 WT RBD protein was coated into 96-well plates at 4 °C overnight. The plates were washed with PBST buffer and blocked with 5% skim milk and 2% bovine albumin in PBS at RT for 1 h. Human ACE2 (Sino Biological, 10108-H08H) or VacW-209 coupled with HRP (Abcam) were mixed with serially diluted mAbs, added into the plates, and then incubated at 37 °C for 1 h. The TMB substrate (Sangon Biotech) was added and incubated at RT for 20 mins and the reaction was stopped by 2M H_2_SO_4_. The readout was detected at a wave length of 450nm. The tested mAbs were 3-fold serially diluted from 10 μg/mL. VRC01 is a HIV-1-specific antibody and used here as a negative control, which is considered as no competition with any SARS-CoV-2-specific mAbs. The percentage of competition was calculated by the formula: (1-OD_450_ of tested mAbs/OD_450_ of VRC01 control) × 100%, and 50% was set as the cutoff indicating an obvious competition.

### Cryo-EM sample preparation and data collection

Aliquots (3 μL) of 3.5 mg/mL mixtures of purified SARS-CoV-2 WT or variant or SARS-CoV spike proteins in complex with excess Fab fragments of VacW-209 were incubated in 0.01% (v/v) Digitonin (Sigma) and then loaded onto glow-discharged (60 s at 20 mA) holey carbon Quantifoil grids (R1.2/1.3, 200 mesh, Quantifoil Micro Tools) using a Vitrobot Mark IV (ThermoFisher Scientific) at 100% humidity and 4 °C. Data were acquired using the SerialEM software on an FEI Tecnai F30 transmission electron microscope (ThermoFisher Scientific) operated at 300 kV and equipped with a Gatan K3 direct detector. Images were recorded in the 36-frame movie mode at a nominal 39,000× magnification at super-resolution mode with a pixel size of 0.339 Å. The total electron dose was set to 60 e^−^ Å^−2^ and the exposure time was 4.5 s.

### Cryo-EM data processing

Drift and beam-induced motion correction was performed with MotionCor2(*6*) to produce a micrograph from each movie. Contrast transfer function (CTF) fitting and phase-shift estimation were conducted with Gctf(*7*). Micrographs with astigmatism, obvious drift, or contamination were discarded before reconstruction. The following reconstruction procedures were performed by using Cryosparc V3(*8*). In brief, particles were automatically picked by using the “Blob picker” or “Template picker”. Several rounds of reference-free 2D classifications were performed and the selected good particles were then subjected to ab-initio reconstruction, heterogeneous refinement and final non-uniform refinement. The resolution of all density maps was determined by the gold-standard Fourier shell correlation curve, with a cutoff of 0.143(*9*). Local map resolution was estimated with ResMap(*10*).

### Cryo-EM model building and analysis

The initial model of nAbs were generated from homology modeling by Accelrys Discovery Studio software (available from: URL: https://www.3dsbiovia.com). The structure of RBD from the structure of WT trimeric spike (6VSB(*11*)) was used as the initial modes of our WT-RBD and Omicron RBD. We initially fitted the templates into the corresponding final cryo-EM maps using Chimera(*12*), and further corrected and adjusted them manually by real-space refinement in Coot(*13*). The resulting models were then refined with phenix.real_space_refine in PHENIX(*14*). These operations were executed iteratively until the problematic regions, Ramachandran outliers, and poor rotamers were either eliminated or moved to favored regions. The final atomic models were validated with Molprobity(*15, 16*). All figures were generated with Chimera or ChimeraX(*17, 18*).

---

**Figure S1.**
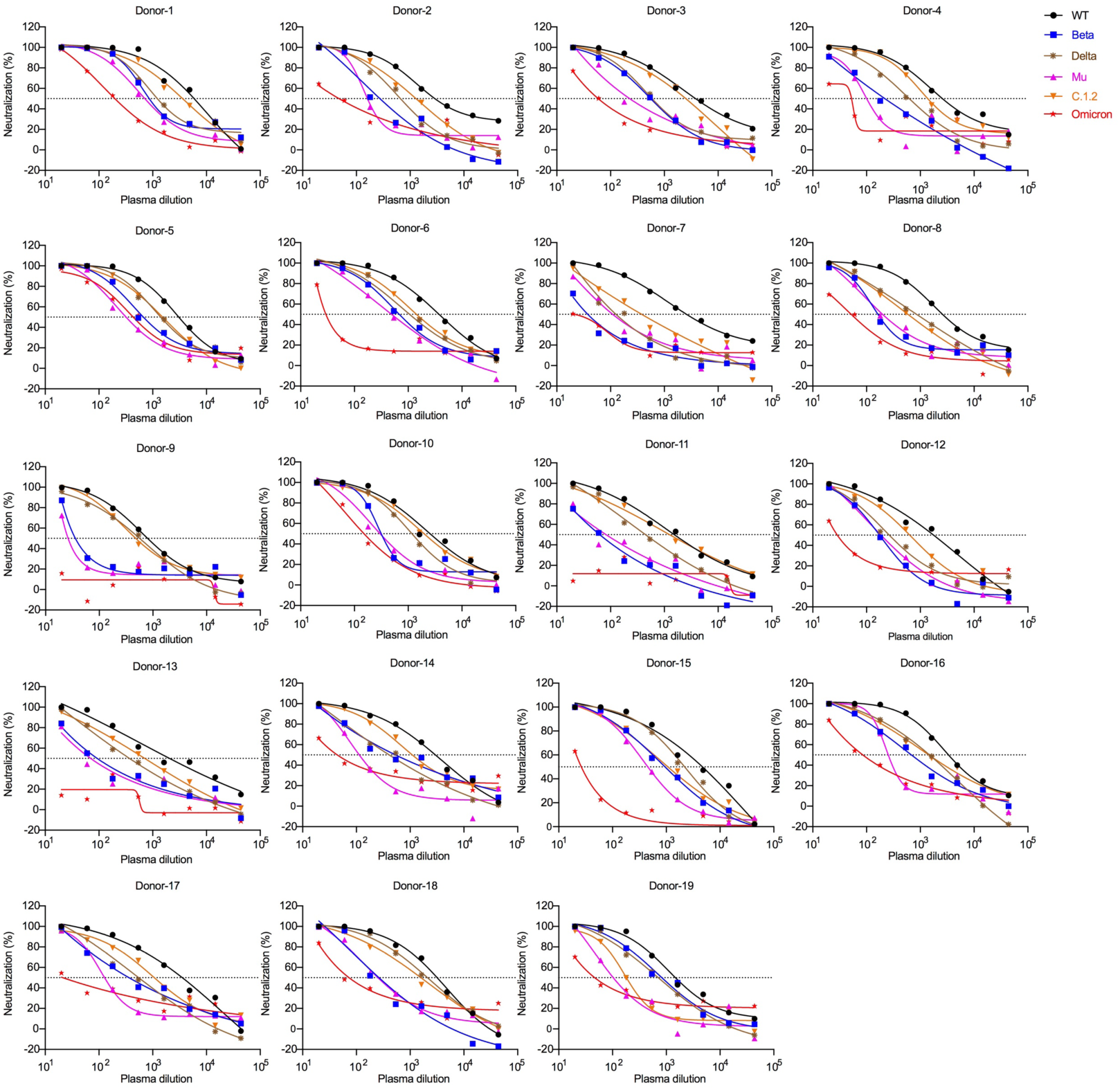
Neutralization curves of plasma samples from 19 convalescent patients recovered from the first wave of COVID-19 pandemic against WT, Beta, Delta, Mu, C.1.2, and Omicron pseudoviruses. All plasma samples were serially 3-fold diluted from 1:20. A 50% reduction in viral infectivity was indicated by a horizontal dashed line. One out of two independent experiments with similar results.

**Figure S2.**
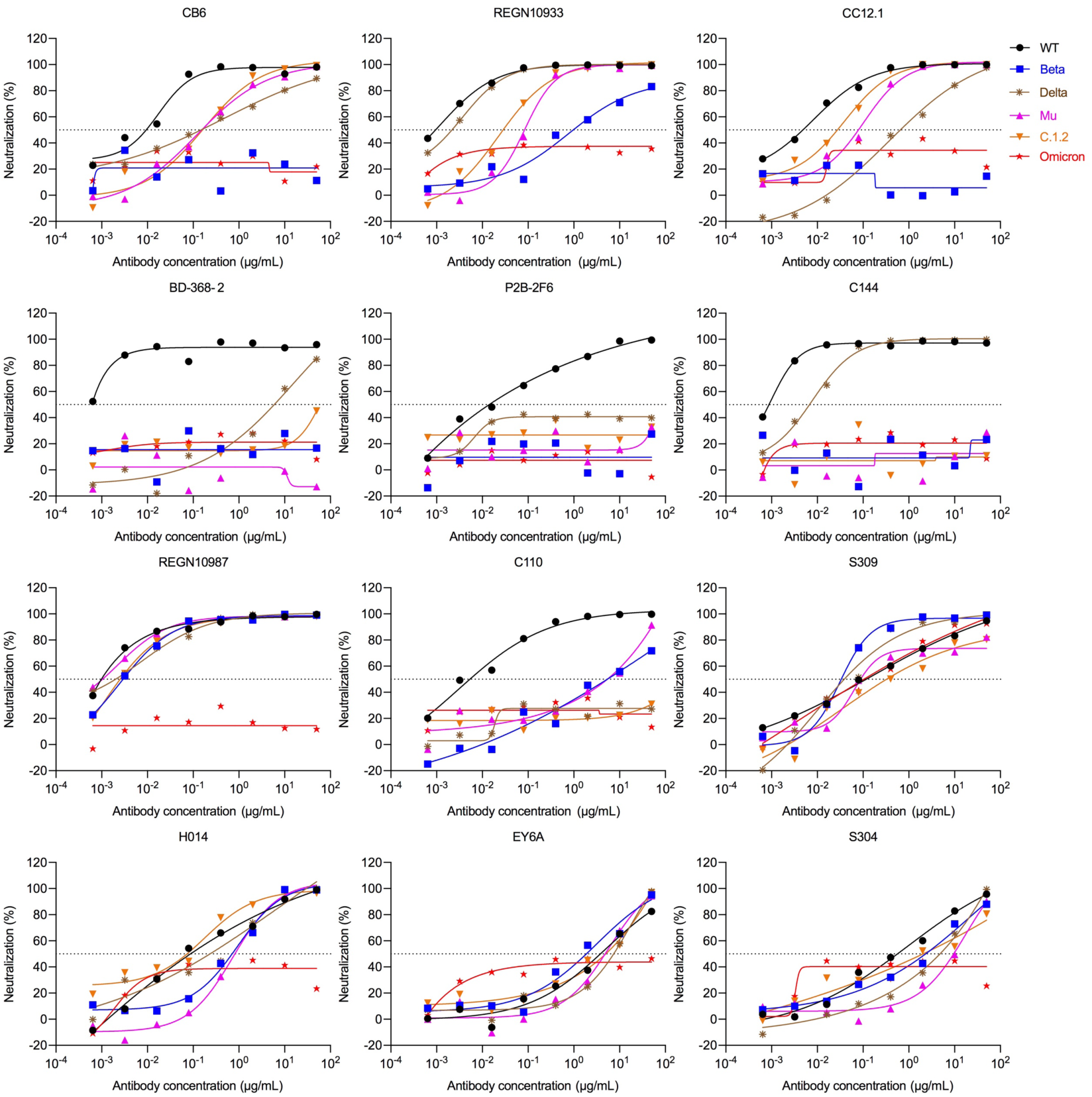
Neutralization curves of 12 representative nAbs against WT, Beta, Delta, Mu, C.1.2, and Omicron pseudoviruses. All nAbs were serially 5-fold diluted from 50 μg/mL. Class 1: CB6, REGN10933, CC12.1; Class 2: BD-368-2, P2B-2F6, C144; Class 3: REGN10987, C110, S309; Class 4: H014, EY6A, S304. A 50% reduction in viral infectivity was indicated by a horizontal dashed line. One out of two independent experiments with similar results.

**Figure S3.**
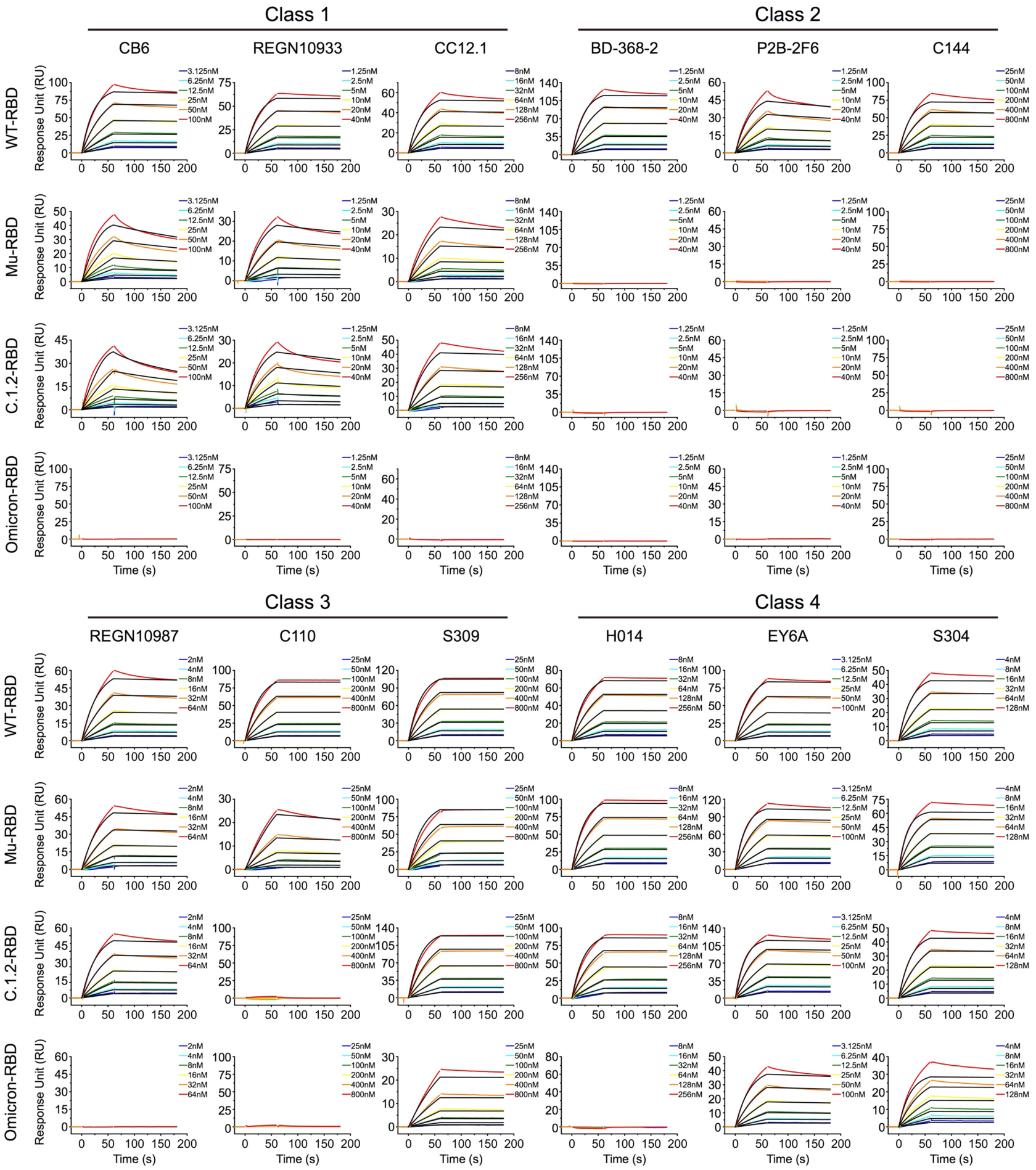
The binding affinities of 12 representative nAbs to SARS-CoV-2 WT, Mu, C.1.2, and Omicron RBD proteins measured by SPR. One out of two independent experiments with similar results.

**Figure S4.**
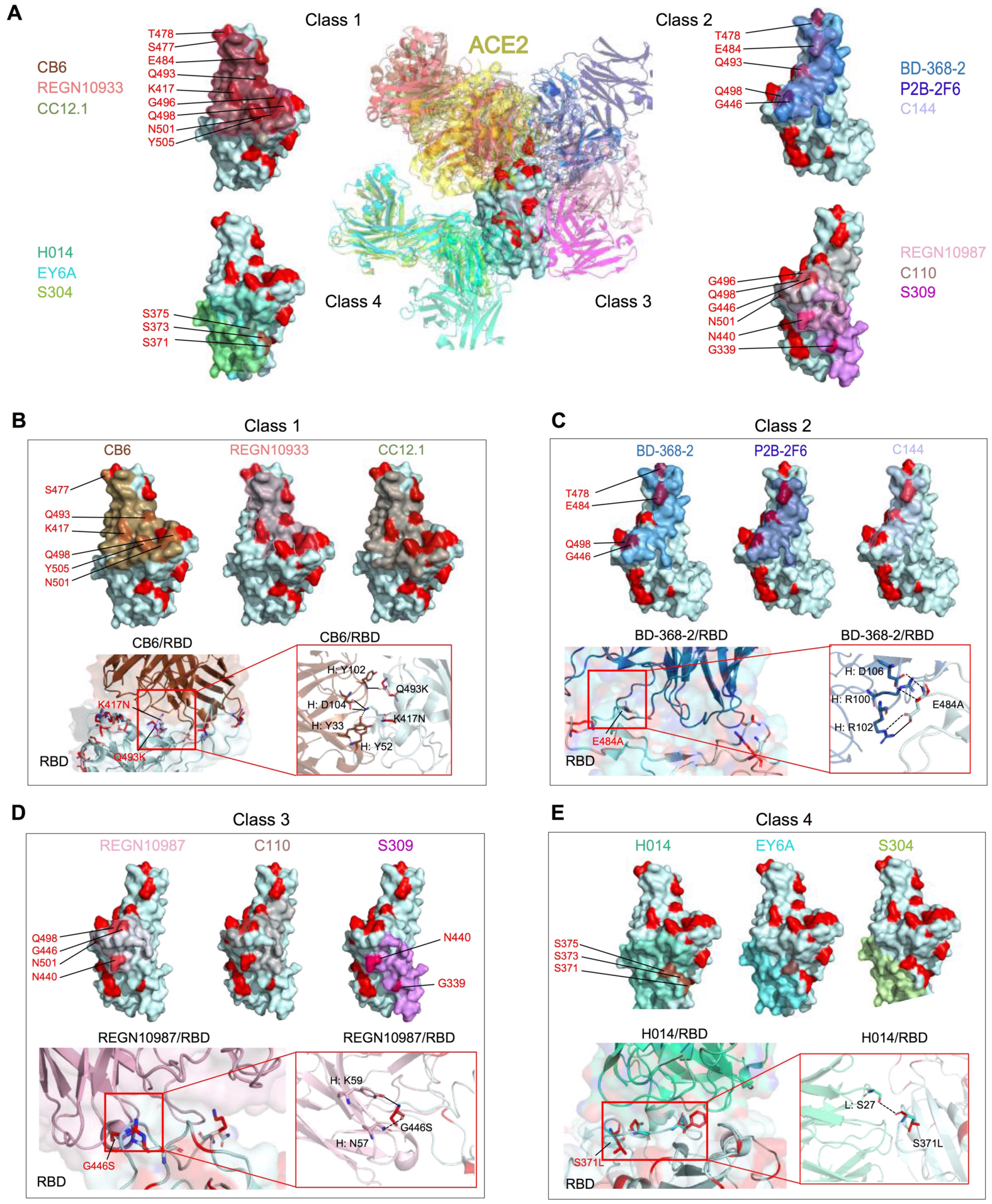
Structural analysis of 12 representative nAbs binding to SARS-CoV-2 RBD. **(A)** Overall structure of ACE2 (PDB: 7DMU) and 12 representative nAbs in complex with SARS-CoV-2 RBD. Footprints of four classes of representative nAbs were represented on the RBD in different colors. The mutated residues appeared in their epitopes were shown in red and labeled beside. The structural analysis of Omicron escaping from nAbs of Class 1 **(B)**, Class 2 **(C)**, Class 3 **(D)**, and Class 4 **(E)**. Cartoon diagrams showing the detailed interface between nAbs and RBD. Hydrogen bond and salt bridge interactions were represented by dashed and black lines, respectively. “H:” indicated antibody heavy chain. “L:” indicated antibody light chain.

**Figure S5.**
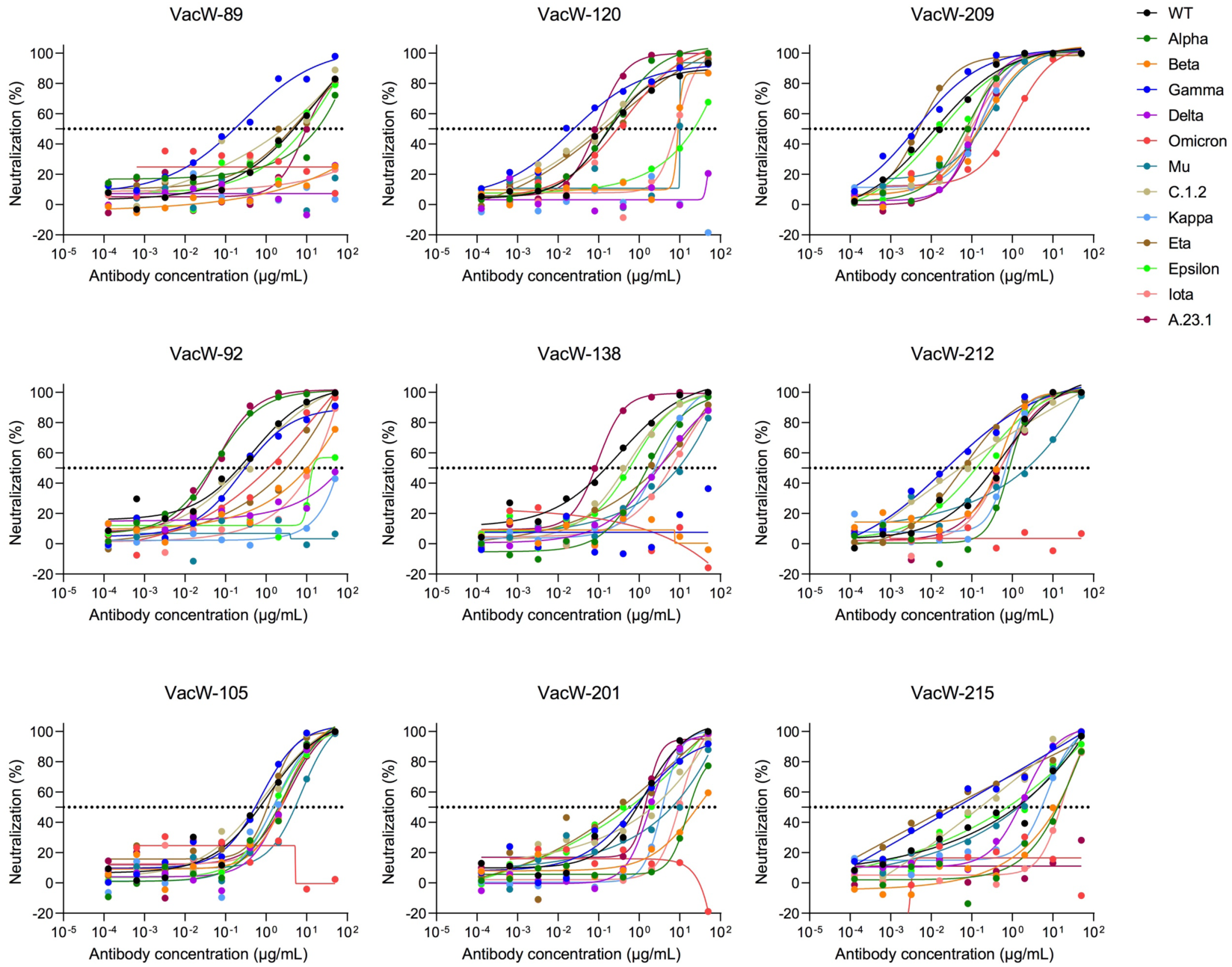
Neutralization curves of 9 nAbs isolated from 2 individuals who received SARS-CoV-2 inactivated vaccines against WT, Alpha, Beta, Delta, Mu, C.1.2, Omicron, and other mutated pseudoviruses. All nAbs were serially 5-fold diluted from 50 μg/mL. A 50% reduction in viral infectivity was indicated by a horizontal dashed line. One out of at least two independent experiments with similar results.

**Figure S6.**
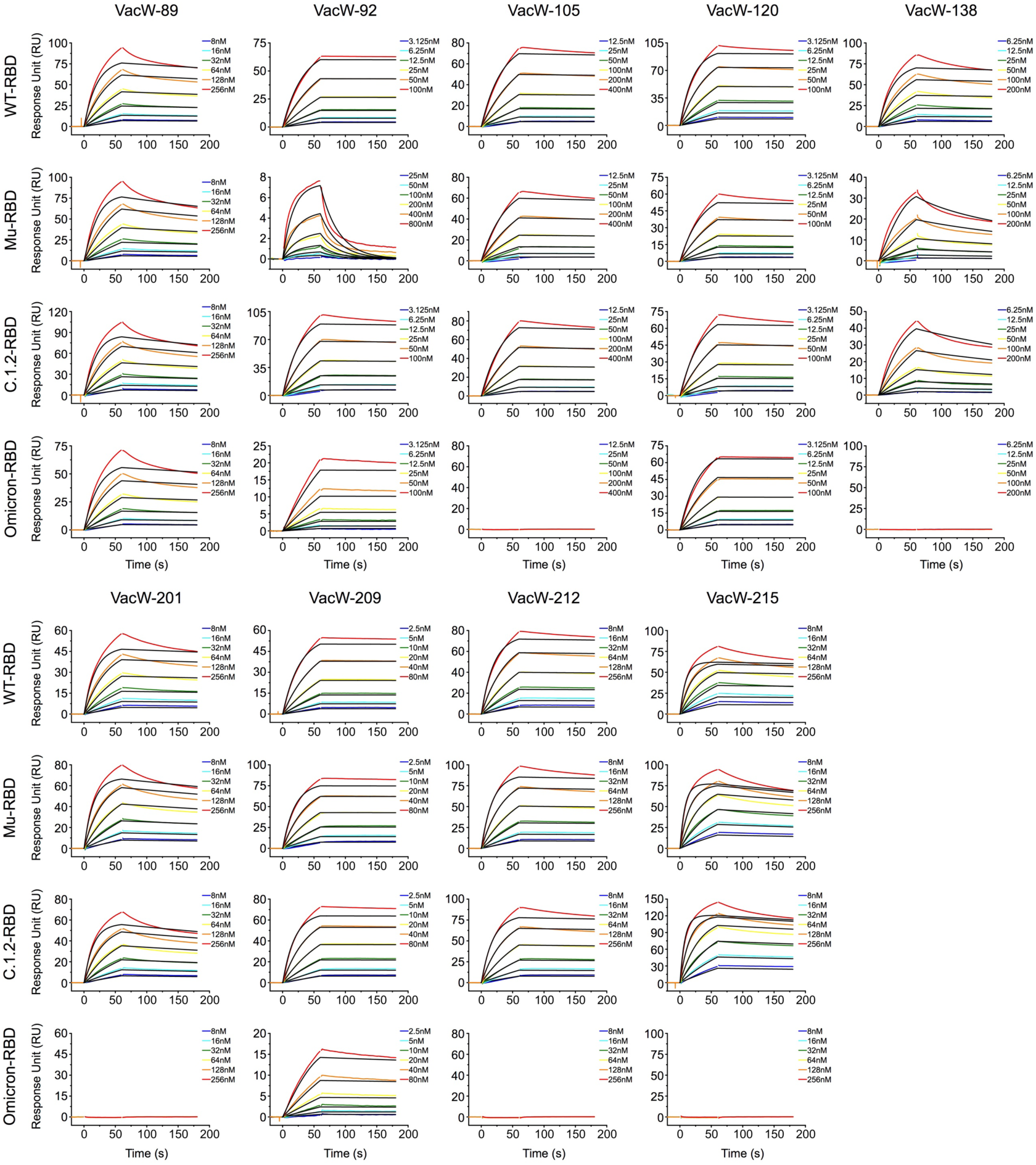
The binding affinities of 9 nAbs isolated from 2 individuals who received SARS-CoV-2 inactivated vaccines to SARS-CoV-2 WT, Mu, C.1.2, and Omicron RBD proteins measured by SPR. One out of two independent experiments with similar results.

**Figure S7.**
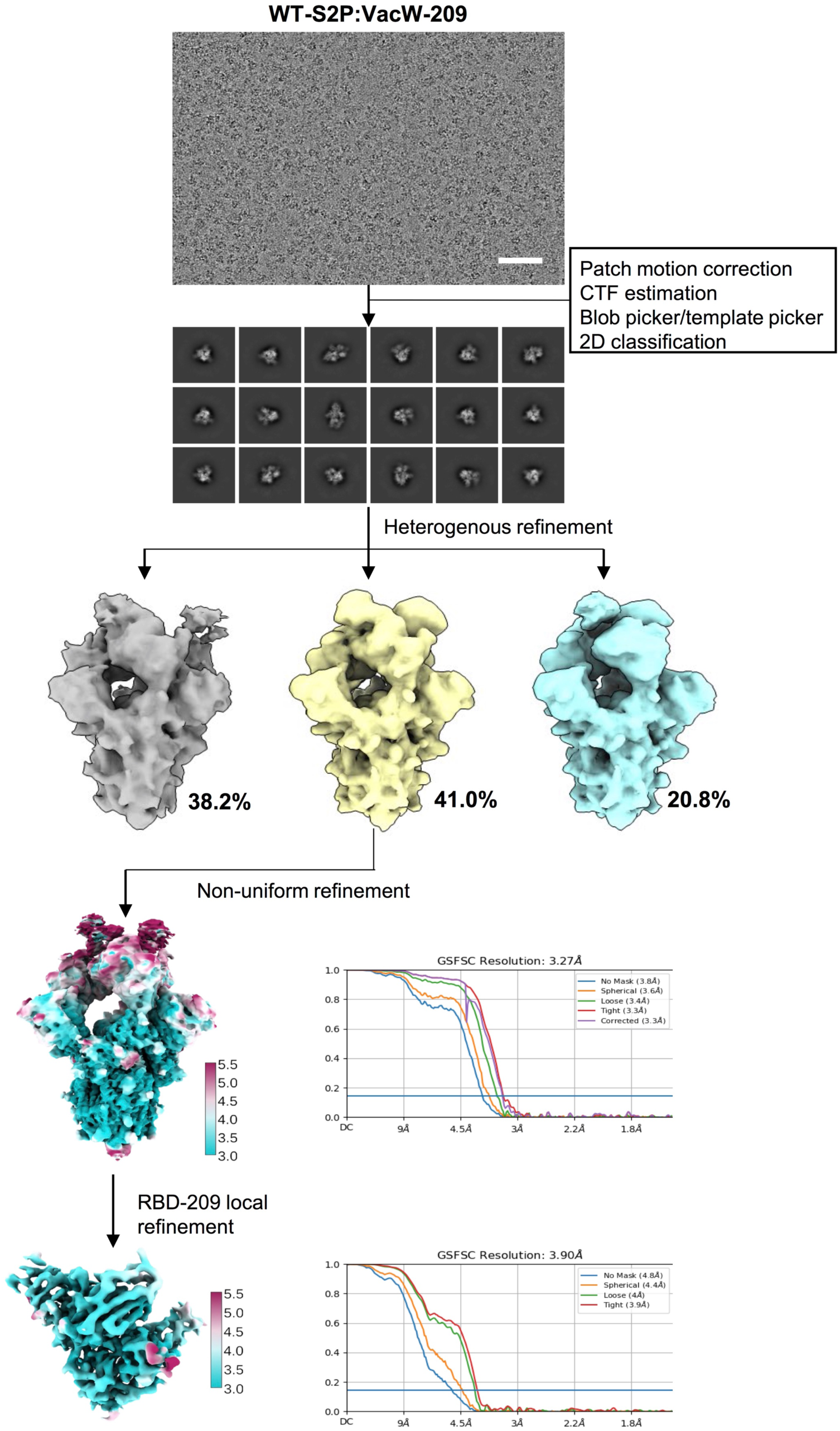
Single-particle cryo-EM images processing workflow and the global and local resolution estimation for the immune complex of SARS-CoV-2 WT-S2P:VacW-209.

**Figure S8.**
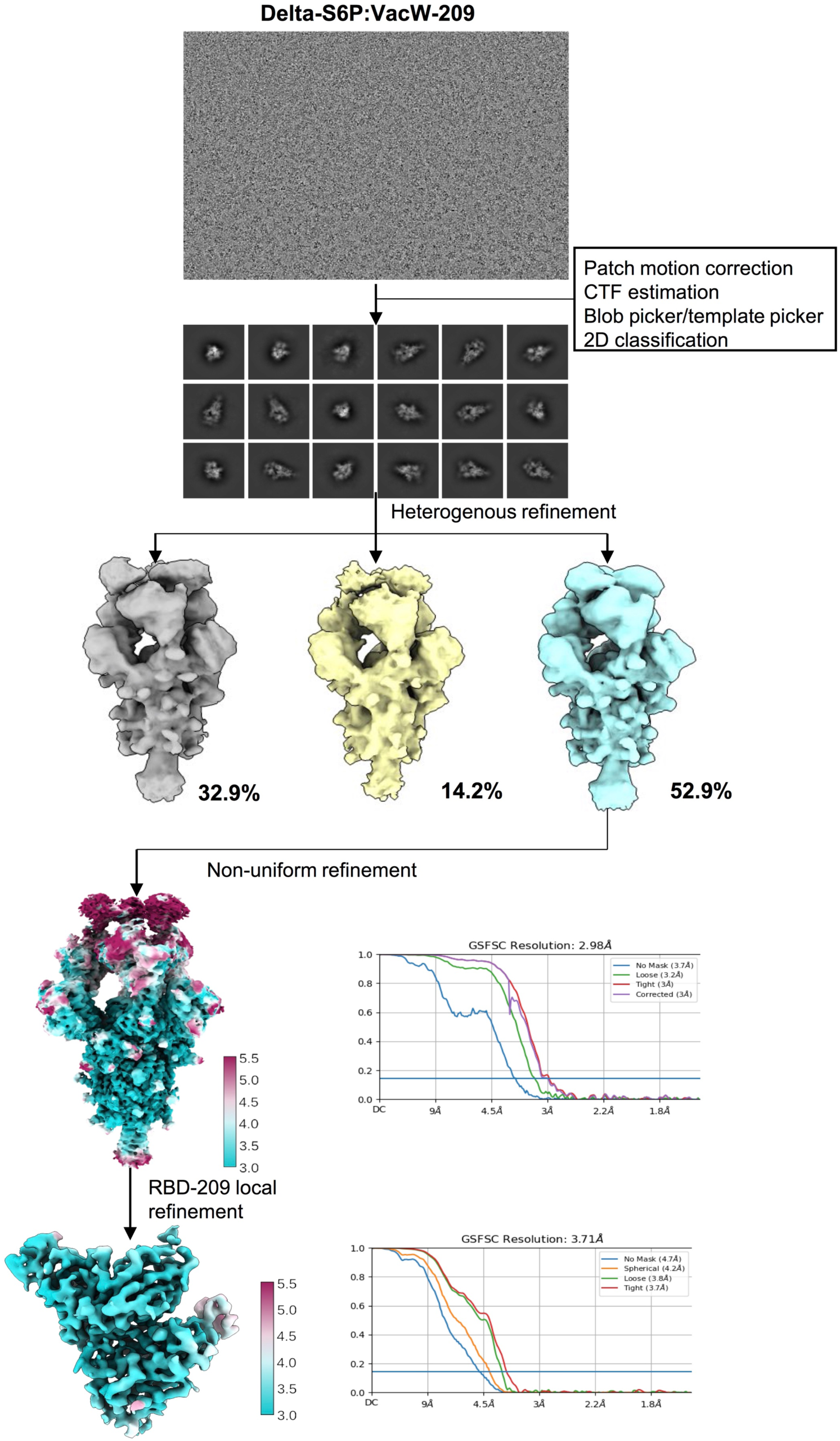
Single-particle cryo-EM images processing workflow and the global and local resolution estimation for the immune complex of SARS-CoV-2 Delta-S6P:VacW-209.

**Figure S9.**
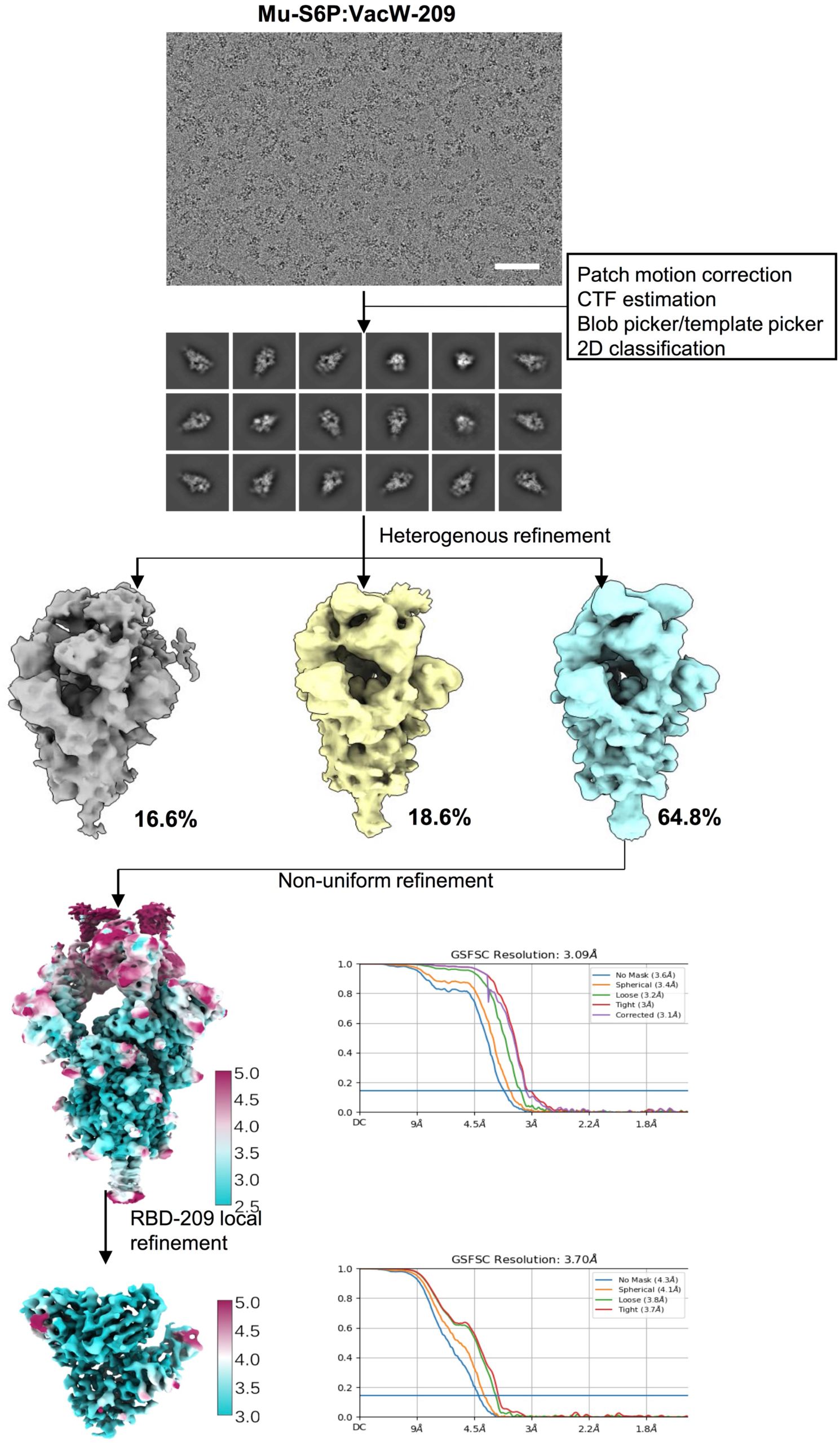
Single-particle cryo-EM images processing workflow and the global and local resolution estimation for the immune complex of SARS-CoV-2 Mu-S6P:VacW-209.

**Figure S10.**
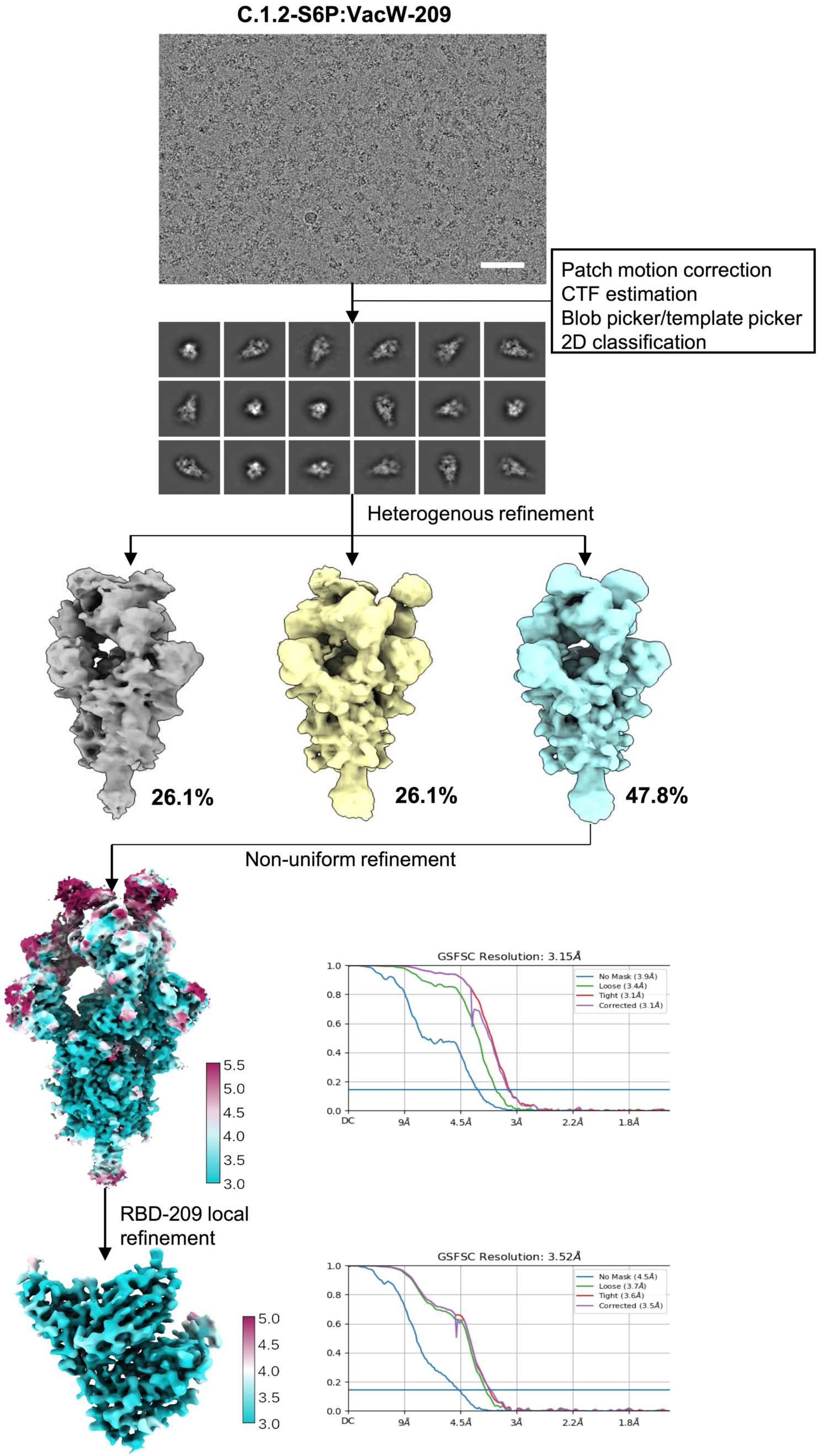
Single-particle cryo-EM images processing workflow and the global and local resolution estimation for the immune complex of SARS-CoV-2 C.1.2-S6P:VacW-209.

**Figure S11.**
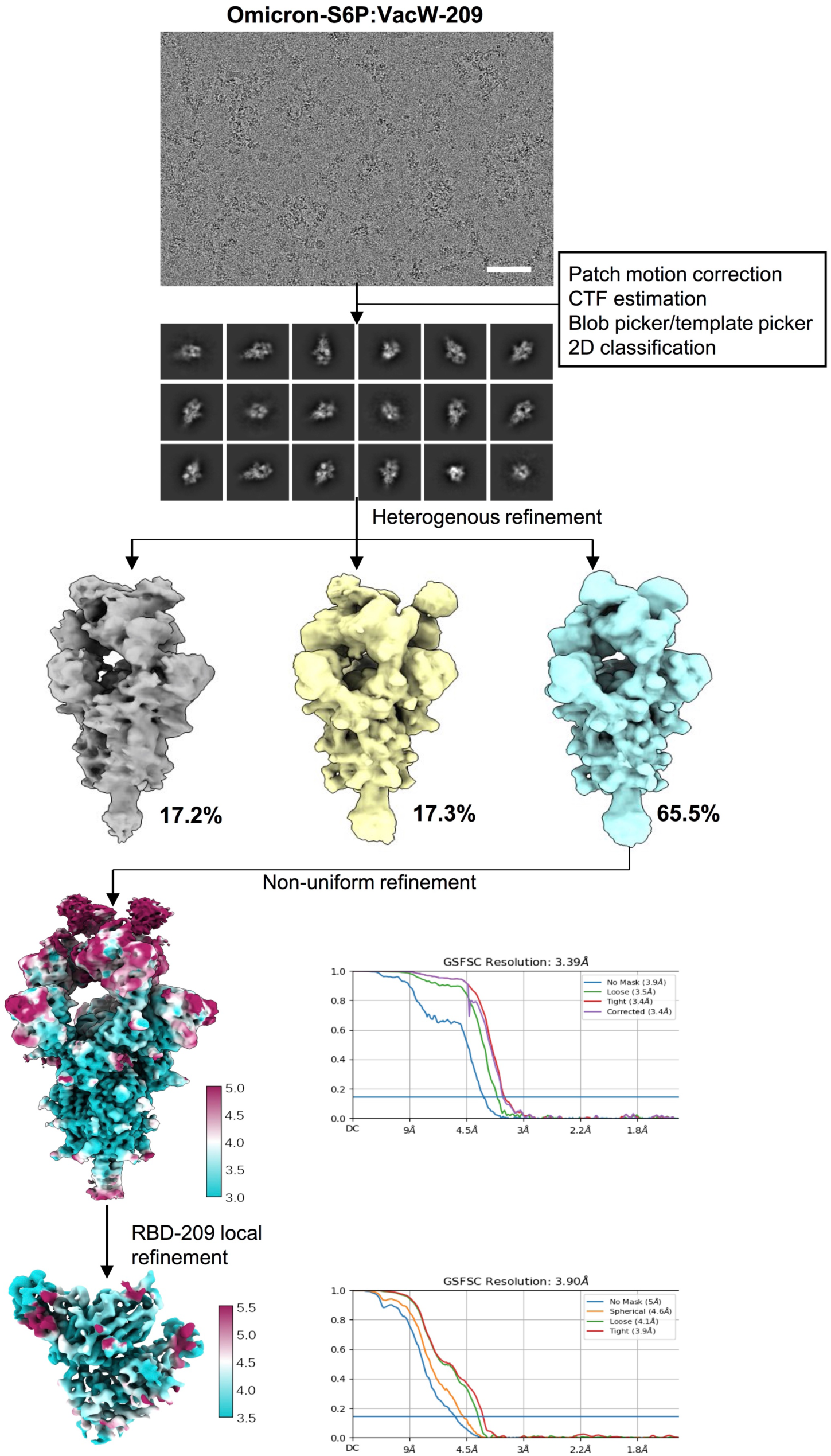
Single-particle cryo-EM images processing workflow and the global and local resolution estimation for the immune complex of SARS-CoV-2 Omicron-S6P:VacW-209.

**Figure S12.**
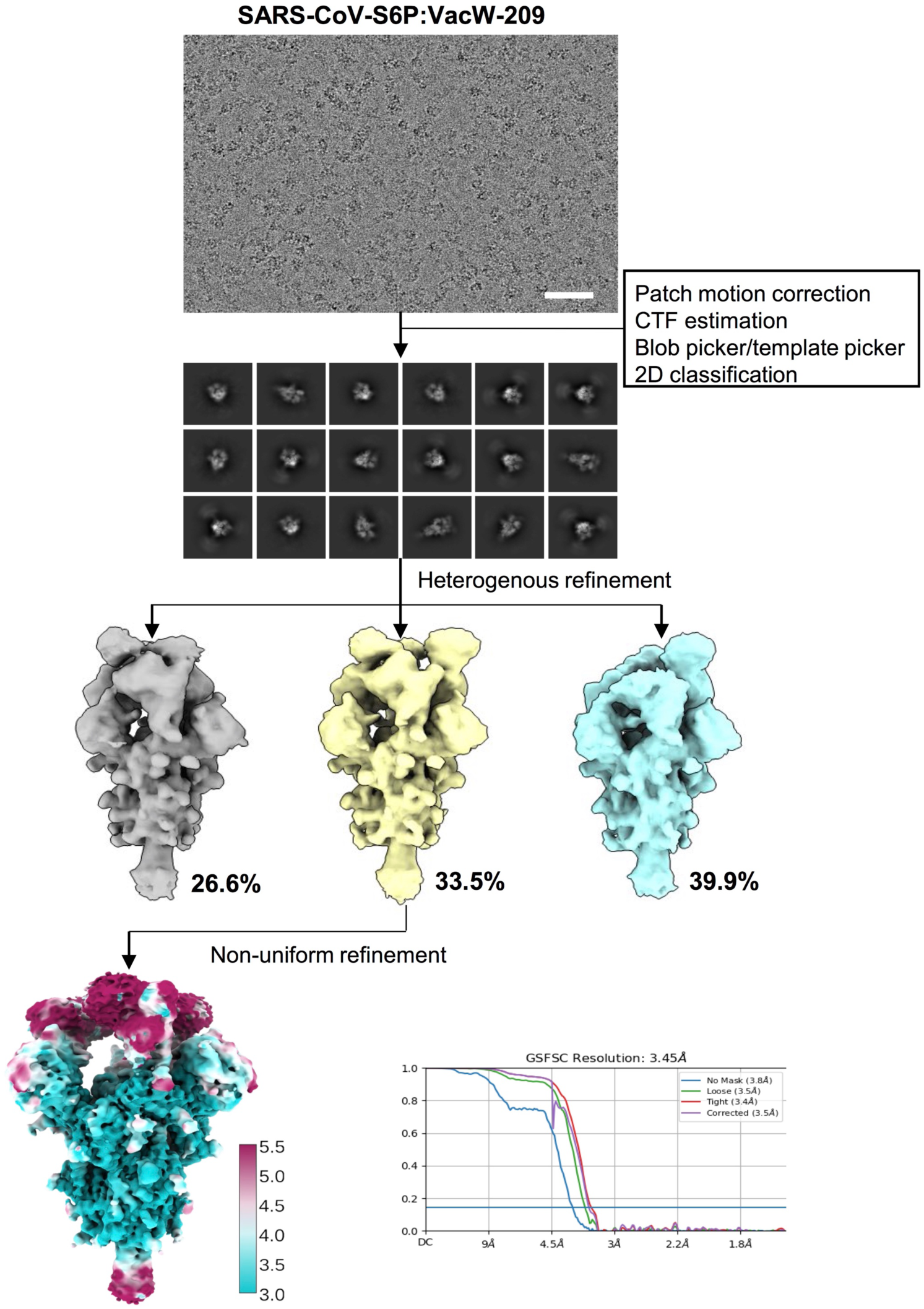
Single-particle cryo-EM images processing workflow and the global and local resolution estimation for the immune complex of SARS-CoV-S6P:VacW-209.

**Figure S13.**
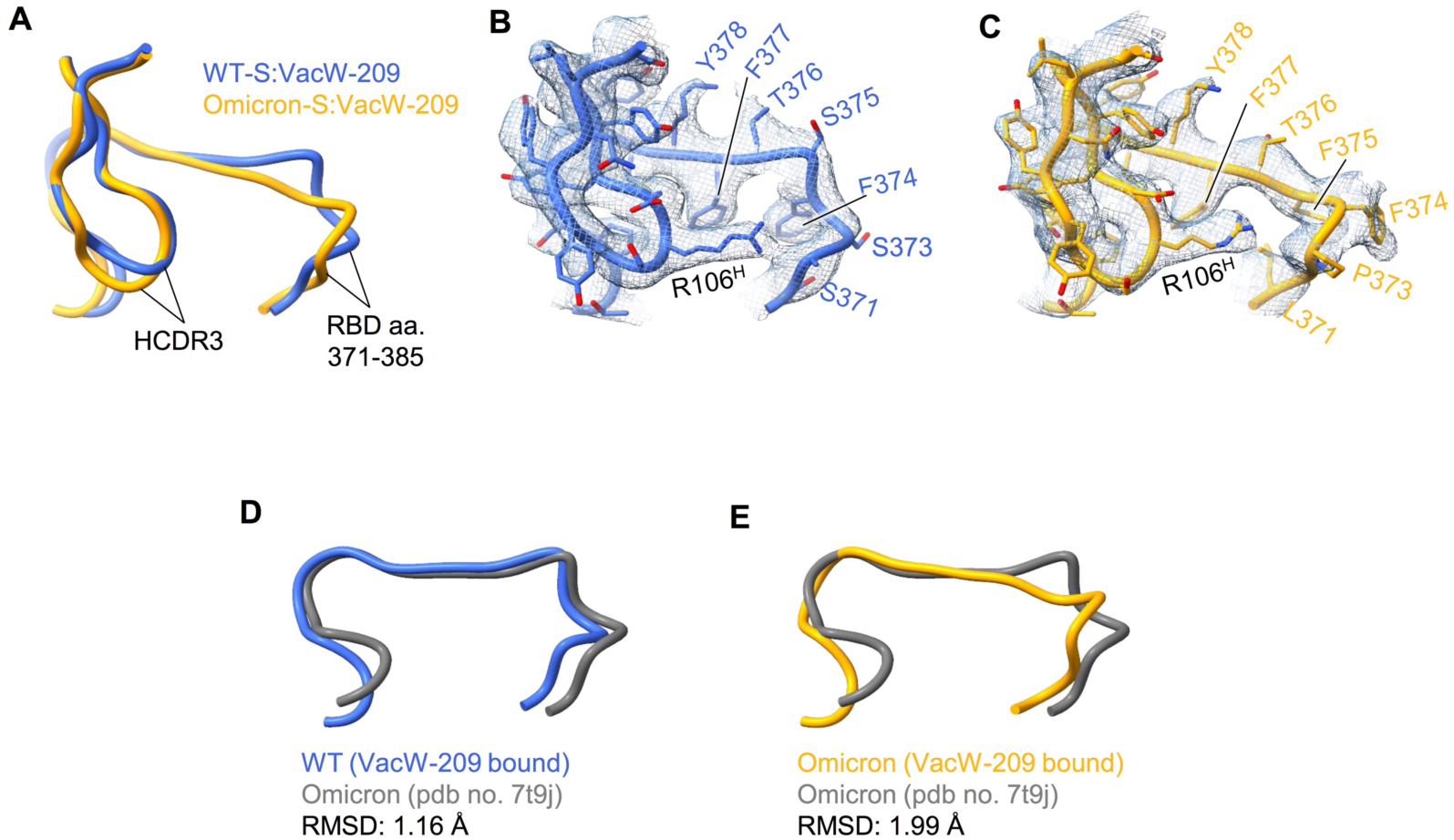
Structural comparison of the interactions of VacW-209 HCDR3 to WT or Omicron RBD. **(A-C)** The HCDR3 (aa. 104-119) and interacted RBD loop (aa. 371-385) from structures of WT-S:VacW-209 (blue) and Omicron-S:VacW-209 (orange) were superimposed shown (A) or separated shown (B-C) with corresponding density maps. **(D-E)** Comparison of RBD loop (aa. 371-385) from WT-S:VacW-209 (D) or Omicron-S:VacW-209 (E) to that from reported crystal structure of Omicron RBD (gray).

**Figure S14.**
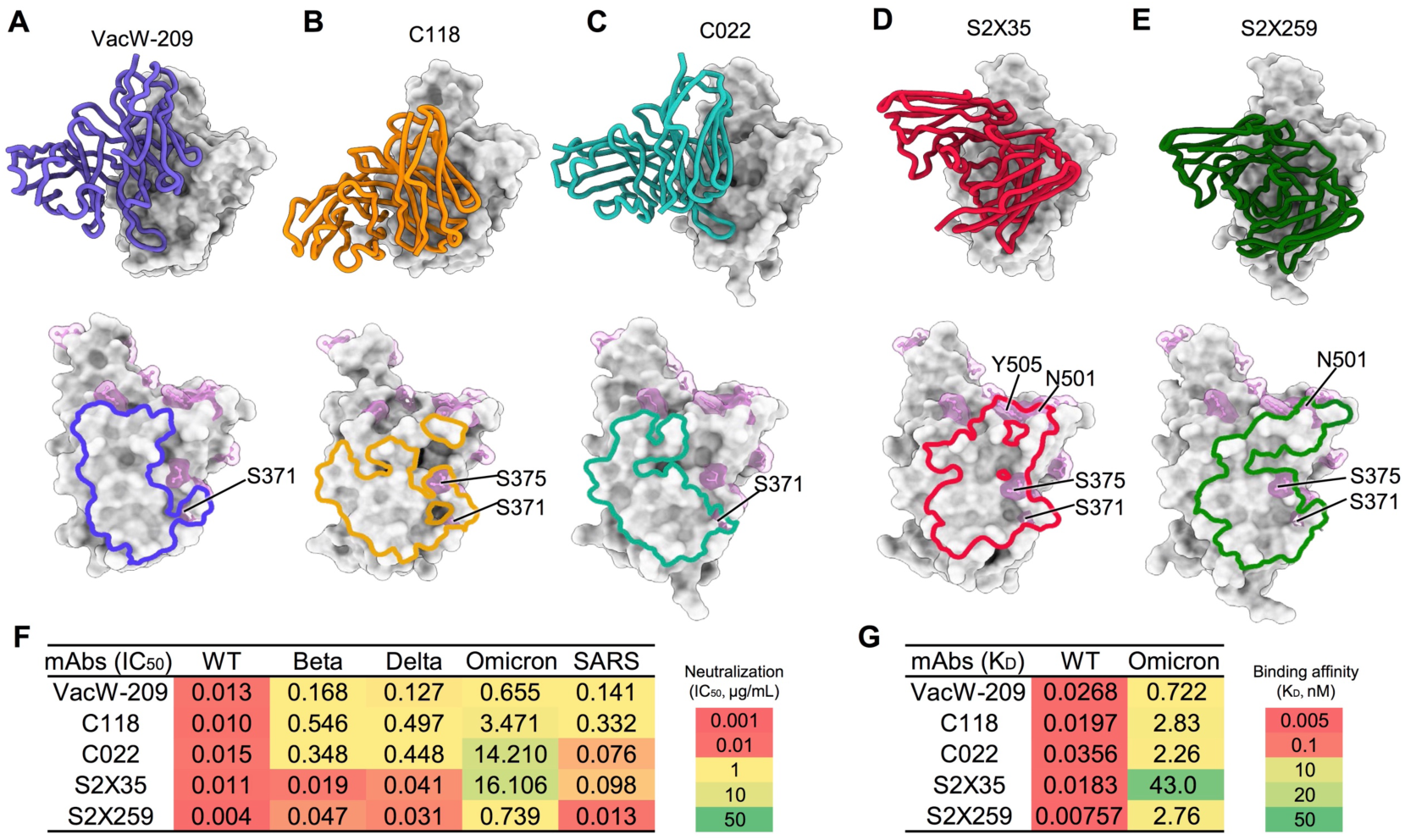
Structural comparison of VacW-209-like nAbs. **(A-E)** Binding modes (upper) and footprints (lower) of VacW-209 (A), C118 (B) (7RKS), C022 (C) (7RKU), S2X35 (D) (7R6W), and S2X259 (E) (7M7W). RBDs are shown as gray surface and nAbs are presented as colored cartoon. The footprints of nAbs and mutations involved in nAbs interactions are labeled in the lower panels. **(F)** The neutralization of VacW-209-like nAbs against SARS-CoV-2 WT, Beta, Delta, Omicron, and SARS-CoV pseudoviruses. **(G)** The binding affinity of VacW-209- like nAbs to SARS-CoV-2 WT and Omicron RBDs by SPR. The data represented here was mean of at least two independent experiments.

**Figure S15.**
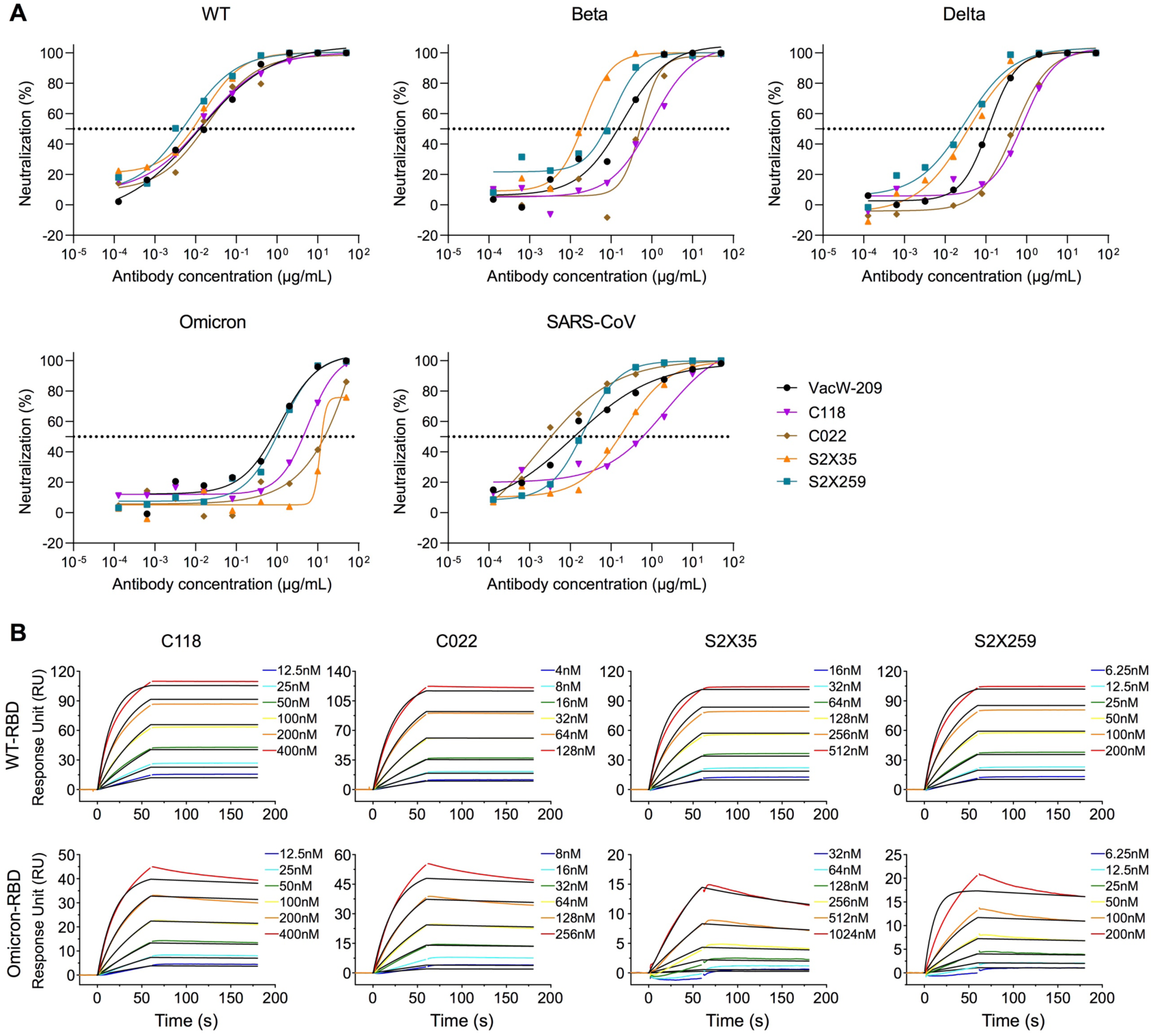
Neutralization (A) and binding affinity (B) curves of VacW-209- like nAbs to SARS-CoV-2 WT, variants, and SARS-CoV. One out of at least two independent experiments with similar results.

**Table S1.** Cryo-EM data collection, refinement and validation statistics of single-antibody Immune-complexes.

|  | SARS-CoV-2-S:209 | SARS-CoV-2-S:209-interface | Delta-S:209 | Delta-S:209-interface | Mu-S:209 | Mu-S:209-interface | C.1.2-S:209 | C.1.2-S:209-interface | Omicron-S:209 | Omicron-S:209-interface | SARS-CoV-S:209 |
| --- | --- | --- | --- | --- | --- | --- | --- | --- | --- | --- | --- |
| **Data collection and processing** |  |  |  |  |  |  |  |  |  |  |  |
| Microscope | FEI TF30 | FEI TF30 | FEI TF30 | FEI TF30 | FEI TF30 | FEI TF30 | FEI TF30 | FEI TF30 | FEI TF30 | FEI TF30 | FEI TF30 |
| Camera | K3 | K3 | K3 | K3 | K3 | K3 | K3 | K3 | K3 | K3 | K3 |
| Magnification | 39,000 | 39,000 | 39,000 | 39,000 | 39,000 | 39,000 | 39,000 | 39,000 | 39,000 | 39,000 | # 39,000 |
| Voltage (kV) | 300 | 300 | 300 | 300 | 300 | 300 | 300 | 300 | 300 | 300 | 300 |
| Electron exposure (e-/Å<sup>2</sup>) | 60 | 60 | 60 | 60 | 60 | 60 | 60 | 60 | 60 | 60 | 60 |
| Defocus range (µm) | 1.2-3.5 | 1.2-3.5 | 1.2-3.6 | 1.2-3.6 | 1.0-3.0 | 1.0-3.0 | 1.0-3.0 | 1.0-3.0 | 1.0-3.2 | 1.0-3.2 | 1.2-3.3 |
| Pixel size (Å) | 0.778 | 0.778 | 0.778 | 0.778 | 0.778 | 0.778 | 0.778 | 0.778 | 0.778 | 0.778 | 0.778 |
| Micrographs (total) | 5342 | 5342 | 2142 | 2142 | 8447 | 8447 | 4062 | 4062 | 8608 | 8608 | 3700 |
| Micrographs (used) | 5098 | 5098 | 2032 | 2032 | 7524 | 7524 | 3703 | 3703 | 7322 | 7322 | 3196 |
| Final particle images (no.) | 320,451 | 336,066 | 129,767 | 389,301 | 323,168 | 969,504 | 225,574 | 225,574 | 150,534 | 451,606 | 271,929 |
| Symmetry imposed | C3 | C1 | C3 | C1 | C3 | C1 | C1 | C1 | C1 | C1 | C1 |
| Map resolution (Å) | 3.27 | 3.90 | 2.98 | 3.71 | 3.09 | 3.77 | 3.15 | 3.52 | 3.39 | 3.90 | 3.45 |
| FSC threshold | 0.143 | 0.143 | 0.143 | 0.143 | 0.143 | 0.143 | 0.143 | 0.143 | 0.143 | 0.143 | 0.143 |
| Map sharpening B factor (Å<sup>2</sup>) | -122.4 | -159.9 | -84.0 | -150.8 | -116.3 | -163.1 | -81.9 | -112.3 | -114.7 | -167.2 | -127.1 |
| **Validation** |  |  |  |  |  |  |  |  |  |  |  |
| MolProbity score | / | 1.82 | / | 1.86 | / | 1.80 | / | 1.87 | / | 1.80 | / |
| Clashscore | / | 5.26 | / | 7.10 | / | 4.03 | / | 6.21 | / | 5.05 | / |
| Poor rotamers (%) | / | 0.28 | / | 0.29 | / | 0.00 | / | 0.57 | / | 0.56 | / |
| RMS (bonds) | / | 0.0063 | / | 0.0080 | / | 0.0060 | / | 0.0127 | / | 0.0055 | / |
| RMS (angles) | / | 1.23 | / | 0.92 | / | 1.30 | / | 1.43 | / | 1.23 | / |
| Ramachdran plot |  |  |  |  |  |  |  |  |  |  |  |
| Favored (%) | / | 90.21 | / | 92.48 | / | 87.53 | / | 90.87 | / | 90.45 | / |
| Allowed (%) | / | 9.79 | / | 7.52 | / | 12.23 | / | 9.13 | / | 9.31 | / |
| Disallowed (%) | / | 0.00 | / | 0.00 | / | 0.24 | / | 0.00 | / | 0.24 | / |

